# Spatial proteomics reveals subcellular reorganization in human keratinocytes exposed to UVA light

**DOI:** 10.1101/2021.09.01.458617

**Authors:** Hellen Paula Valerio, Felipe Gustavo Ravagnani, Angela Paola Yaya Candela, Bruna Dias Carvalho da Costa, Graziella Eliza Ronsein, Paolo Di Mascio

**Affiliations:** Department of Biochemistry, Institute of Chemistry, University of São Paulo, São Paulo, 05508-000, Brazil

## Abstract

The effects of UV light on the skin have been extensively investigated. However, systematic information about how exposure to UVA light, the least energetic but the most abundant UV radiation reaching the Earth, shapes the subcellular organization of proteins is lacking. Using subcellular fractionation, mass spectrometry-based proteomics, machine learning algorithms, immunofluorescence, and functional assays, we mapped the subcellular reorganization of the proteome of human keratinocytes in response to UVA light. Our workflow quantified and assigned subcellular localization for over 1600 proteins, of which about 200 were found to redistribute upon UVA exposure. Reorganization of the proteome affected modulators of signaling pathways, cellular metabolism, and DNA damage response. Strikingly, mitochondria were identified as one of the main targets of UVA-induced stress. Further investigation demonstrated that UVA induces mitochondrial fragmentation, up-regulates redox-responsive proteins and attenuates respiratory rates. These observations emphasize the role of this radiation as a potent metabolic stressor in the skin.

## Introduction

Ultraviolet-A (UVA) light (315–400 nm) constitutes about 95% of all ultraviolet radiation (UVR) that reaches the Earth (Schuch et al., 2017). The causal association between UVR exposure and skin cancer is well established, but epidemiology has little capacity to distinguish between the carcinogenic effects of UVA and UVB (El Ghissassi et al., 2009). At the molecular level, the effects of UVA and UVB in skin cells are of different natures, suggesting that each wavelength range defines a different path towards malignant transformation (Premi et al., 2015).

For example, UVB is absorbed by pyrimidines, giving rise to cyclobutane pyrimidine dimers (CPD) and pyrimidine (6-4) pyrimidone photoproducts. Thus, UVB’s carcinogenic action depends on the direct generation of mutagenic DNA lesions (Schuch et al., 2017). On the other hand, UVA photons are poorly absorbed by the DNA, being more relevantly absorbed by other cellular chromophores (Ikehata, 2018). In this sense, UVA relies on the generation of photoexcited species, such as singlet oxygen, that may lead to oxidative damage (Wondrak et al., 2006).

Skin cells orchestrate complex responses to light stress, coordinating gene expression, metabolism and protein function (Chen et al., 2014). Protein function is fine-tuned in a sophisticated manner, involving modulations in abundance, chemical modifications, and spatial and temporal delimitations (Thul et al., 2017). Mutational dynamics is the primary driver of carcinogenesis. However, modulation of metabolism and protein function can contribute to this process by impacting signaling, organelle interactions and cell fate decisions towards apoptosis, senescence or malignant transformation (de Gruijl et al., 2001; El Ghissassi et al., 2009). Importantly, besides impacting cell fate decisions, UV-induced changes in cellular physiology can alter differentiation and stem cell division, accelerating the growth of mutant cell clones in the skin. In this sense, changes in cellular physiology may be numerically more important to cancer development than the original oncogenic mutations (Klein et al., 2010).

Even though the effects of UVR on DNA modification (Moreno et al., 2020), gene expression (He et al., 2004), protein expression (Edifizi et al., 2017) and post-translational modifications (Elia et al., 2015; Zhou et al., 2016) have been investigated, information about how specific UVR components shape the subcellular organization of proteins in cells is still lacking. Advances in high-throughput mass spectrometry (Breker and Schuldiner, 2014; Larance and Lamond, 2015) and microscopy (Mattiazzi Usaj et al., 2016; Thul et al., 2017) and machine learning applications for these techniques (Gatto et al., 2014a; Lundberg and Borner, 2019) allow proteome-wide investigations into subcellular localization dynamics and organellar communication in cells under stress. Spatial or organellar proteomics workflows may combine cell fractionation with mass spectrometry to characterize changes in protein levels in multiple subcellular niches (Lundberg and Borner, 2019). Indeed, methods such as Protein Correlation Profiling (PCP) (Andersen et al., 2003; Foster et al., 2006), Hyperplexed Localisation of Organelle Proteins by Isotope Tagging (HyperLOPIT) (Geladaki et al., 2019; Mulvey et al., 2017) and other organellar mapping approaches (Itzhak et al., 2017; Jean Beltran et al., 2016) have been developed to monitor protein dynamics over space in an unbiased manner.

The principle behind these methodologies is to quantify the distribution of proteins across subcellular fractions under different biological conditions. The fractionation profiles of proteins reflect the complexity of subcellular localization better than the presence or absence in a single purified fraction. Thus, they are used as an input for learning algorithms, allowing the classification of subcellular localization. Recently, a machine learning pipeline classified translocation events between subcellular niches by allowing the comparison of fractionation profiles under different biological conditions (Kennedy et al., 2020).

In light of these advances, we used spatial proteomics coupled with machine learning techniques to systematically analyze the subcellular reorganization of the proteome of skin cells in response to UVA radiation. Our results show that a low UVA dose, equivalent to about 20 minutes of midday sun exposure (Halliday and Rana, 2008), leads to spatial remodeling of the skin cells’ proteome. We found that the spatial stress response relies on changes in mitochondrial dynamics, redox modulations and a nucleocytoplasmic translocation triggered by DNA damage. Furthermore, our results provide a resource for further investigations of UVA-triggered protein dynamic events.

## Results

### Workflow used to investigate proteome remodeling of skin cells under UVA light stress

An overview of the experimental protocol is shown in **Figure 1A**. In the experimental pipeline, HaCaT skin cells were exposed to a non-cytotoxic low dose of UVA light (6 J/cm^2^, using a simulator of the solar UVA spectrum) or kept in the dark under the same environmental conditions. Mock-treated and UVA-exposed cells were collected, the plasma membranes were lysed in hypoosmotic solution, and the organelles were separated by differential centrifugation. Fractions were collected after each centrifugation step, and proteins were quantified in each fraction by conventional label-free mass spectrometry. A total of 5351 protein groups were identified and quantified in 90 samples, comprising nine fractions for each of the five biological replicates of each condition. The dataset was filtered for proteins with label-free quantifications (LFQ) greater than zero and in at least 50% of all samples, yielding a matrix of 1650 protein groups. This step was performed to exclude proteins that were irregularly quantified across replicates and fractions. The remaining missing values were imputed. Briefly, for proteins with at least 4 valid values across the 5 replicates of a given fraction, the missing value was imputed as the average of the 4 valid values. The remaining missing values were imputed as the minimum value of the sample. This imputation approach should account for both values missing at random and not at random, which are commonly present in shotgun proteomics datasets (Dabke et al., 2021; Liu and Dongre, 2021). **Figure S1A-D** shows the results obtained during optimization of the imputation method and **Figure S1E-F** shows the final imputed subcellular maps of control and irradiated samples. **Supplementary Spreadsheet S1 and S2** provide the unfiltered LFQ dataset without imputation and the filtered dataset with imputation, respectively.

**Figure 1.**
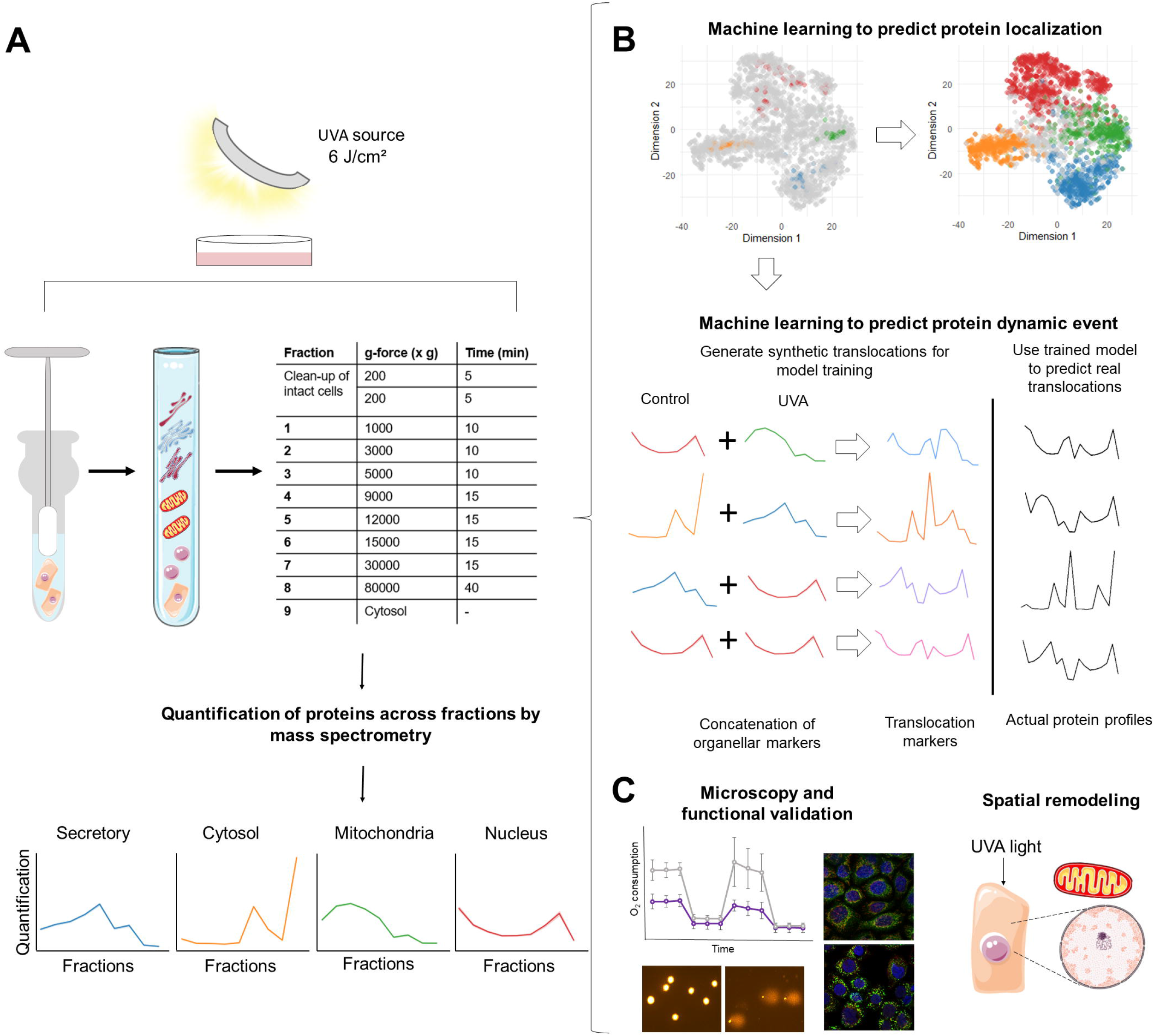
Proteomic approach to define spatial changes in protein distribution upon light stress. **(A)** Experimental protocol. **(B)** Computational pipelines used to define changes in subcellular organization promoted by UVA light in HaCaT cells. **(C)** Validation of results using traditional biochemical assays.

Next, to assess if the dataset’s structure reflected subcellular localization, we used three complementary approaches to inspect data quality, predict subcellular localization and infer protein dynamic events. First, we used t-SNE as a dimensionality reduction method. Proteins in the t-SNE plots were colored according to subcellular localization information available in three different databases (Uniprot, Gene Ontology and Cell Atlas), aiming to inspect clusters formation. Second, we used a neural networks algorithm to assess if subcellular localization could be attributed accurately by learning the fractionation patterns of organellar markers with well-established localization.

Lastly, after validating the dataset’s structure, we used the Translocation Analysis of Spatial Proteomics (TRANSPIRE) computational pipeline (Kennedy et al., 2020), which is based on a gaussian process classifier, to investigate changes in the subcellular landscape induced by UVA light in human keratinocytes. An overview of the computational workflow is presented in **Figure 1B**. Some of the results obtained by TRANSPIRE were further validated by conventional biochemical assays (**Figure 1C**).

### Validating the resolving power of the fractionation method

Following our workflow, we first inspected the t-SNE plot generated from the filtered dataset to reduce dimensionality and detect the presence of clusters. The plot revealed the presence of four main clusters in distinct regions (**Figure S2**). When proteins were colored according to subcellular localization obtained from three different databases (Uniprot, Cell Atlas and Gene Ontology), we found that the four clusters represented four distinct subcellular environments: the nucleus, cytosol, mitochondria, and secretory organelles. The database classifications were binned such that secretory organelles included proteins from the ER, peroxisome, Golgi, lysosome and plasma membrane (**Figure S2**).

Since this analysis showed that the fractionation scheme provides the resolution necessary for differentiating well these four main subcellular compartments, we curated organellar markers for each compartment to investigate if protein localization could be classified based on the fractionation scheme. We obtained a set of organellar markers from curating subcellular localization information available in the Gene Ontology (GO) and Uniprot. We also obtained a few classifications from a previous subcellular proteomics study (Geladaki et al., 2019). For a protein to be considered an organellar marker, it had to be classified in both databases as uniquely pertaining to one subcellular niche among the four compartments (i.e., cytosol, nucleus, mitochondria and secretory) established through dimensionality reduction. Organellar markers also had to be consistently present and quantified in all replicates of our experiment.

Based on these criteria, 550 organellar markers were curated into four subcellular niches: the cytosol (117), nucleus (201), mitochondria (109), and secretory organelles (123). The complete set of organellar markers used in this study can be found in **Supplementary Spreadsheet S3**. Fractionation profiles of markers from different compartments present characteristic shapes, demonstrating that proteins from the same subcellular niche tend to fractionate similarly (profile plots, **Figure 2A**). The t-SNE supports the patterns observed in the profile plots, showing that organellar markers from different compartments cluster in separate plot regions, while markers of the same compartment cluster similarly (**Figure 2B**). These plots also reveal a shift in the mitochondrial cluster in the subcellular map of irradiated cells relative to controls. This shift seems to approximate the mitochondrial and secretory clusters (a more detailed 2D version of this plot, with labels of the markers, is shown in **Figure S1E-F**).

**Figure 2.**
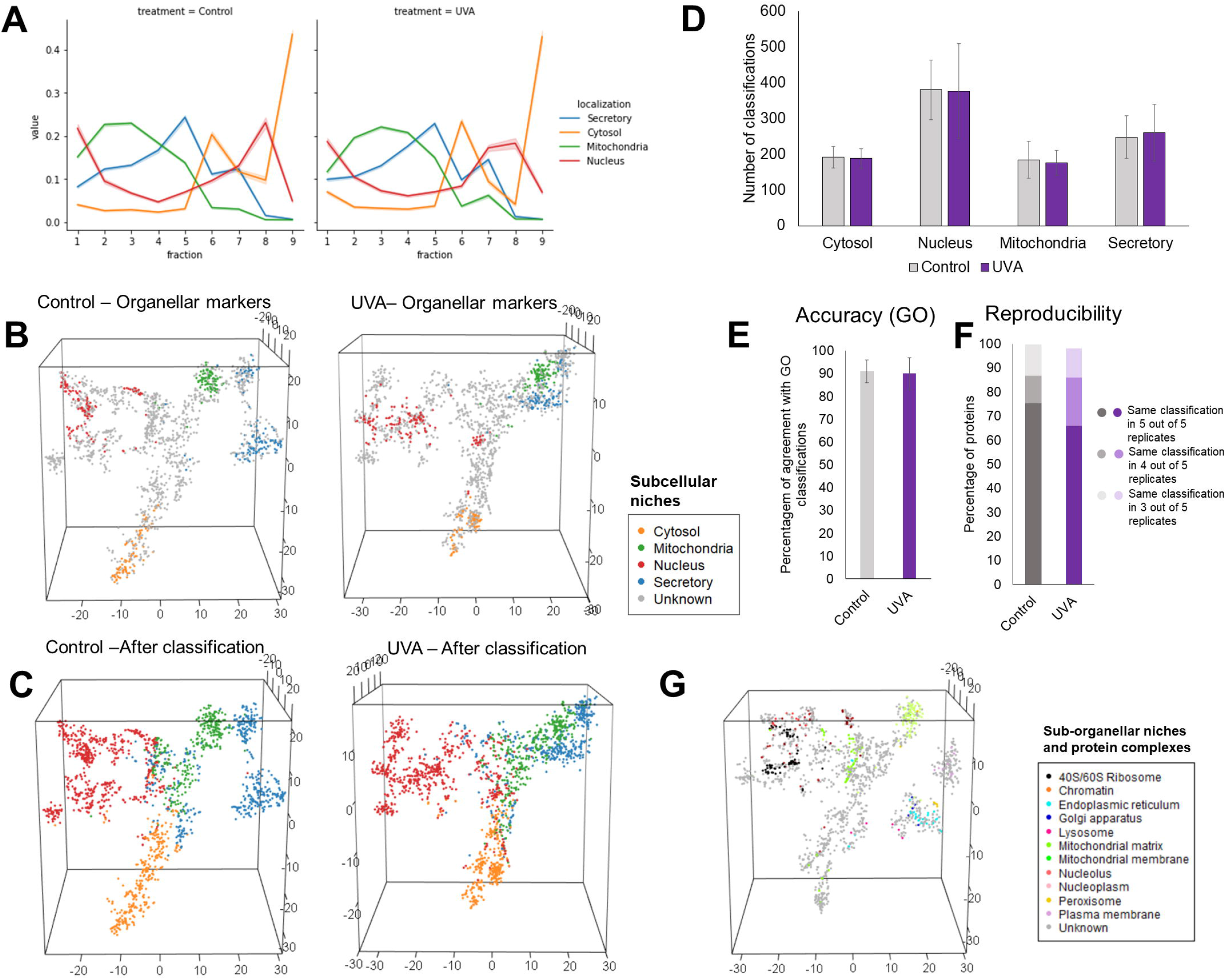
Identification of subcellular patterns in HaCaT’s spatial proteomics dataset. **(A)** Profile plots of organellar markers in the HaCaT dataset. Shadowed intervals represent standard errors, and values represent means. Five biological replicates per group were employed for the spatial proteomics experiment. **(B)** 3D representation of subcellular fractionation data using t-SNE. The maps were colored according to organellar markers. **(C)** t-SNE plots of all control and irradiated samples overlaid with the most frequent classifications obtained for each protein across replicates using the neural network algorithm for predicting localization. **(D)** Numbers of proteins assigned to the cytosol, mitochondria nucleus and secretory organelles by condition. Bars represent the mean number of proteins assigned to each compartment, and error bars represent the standard deviation. **(E)** Accuracy of the neural network predictions obtained by comparing the predicted subcellular localizations with Gene Ontology information. Bars represent means per condition, and error bars represent standard deviations. **(F)** Reproducibility of classifications across replicates. Bars represent the percentage of proteins that received the same classification in 3, 4 or 5 biological replicates out of the total 5. **(G)** t-SNE plots overlaid with sub-organellar markers, colocalizing protein complexes and markers of secretory compartments.

Following this analysis, a neural networks algorithm implemented in pRoloc (Gatto et al., 2014b), utilizing as references of each compartment the fractionation profiles of curated organellar markers, classified proteins into four discrete subcellular compartments. To assess the reproducibility of classifications across replicates, we applied the algorithm to each of the five biological replicates of each condition separately (the F1 scores obtained during hyperparameter optimization of the neural networks are shown in **Figure S3**). The output of the algorithm can be found in **Supplementary Spreadsheet S4**. **Figure 2C** contains the t-SNE plots representing the most frequent classification of each protein across the five replicates for each condition. All of the 1100 protein groups were classified into four subcellular niches: the cytosol (190 ± 30 proteins, considering the mean and standard deviation across replicates), nucleus (379 ± 83), mitochondria (180 ± 43), and secretory organelles (254 ± 70), with slight differences for the total number of classifications between conditions (**Figure 2D**). All classifications obtained from the machine learning algorithm are accompanied by classification probability scores that reflect the reliability of the assignment. In this context, low scores are often associated with profiles not directly modeled by the organellar markers used in the algorithm (e.g., multilocalized proteins) (Jean Beltran et al., 2016). To account for multilocalized proteins, we filtered proteins with low prediction scores (the numbers above represent the filtered classifications). Considering the filterered dataset, the subcellular localization classifications obtained for each replicate were compared to GO classifications. The results revealed that the averaged neural networks algorithm achieved a mean prediction accuracy of 91% in control samples and 90% in treated samples (**Figure 2E**). Classifications were also highly reproducible, with almost 100% of all proteins in the treated and control samples receiving the same classification in at least 3 out of 5 biological replicates (**Figure 2F**).

In addition, we analyzed if the dataset could provide sub-organellar resolution by differentially coloring markers of sub-organellar compartments, protein complexes and secretory organelles in the t-SNE plot. Markers of sub-organellar and secretory compartments were mostly obtained from Geladaki et al., 2019. A few markers were obtained from Uniprot, Mitocarta and Gene Ontology. The results indicate a partial divide between the mitochondrial matrix, membrane and nuclear subniches, such as the nucleolus, nucleoplasm, and chromatin clusters (**Figure 2G**). Moreover, the t-SNE plot also reveals that specific protein complexes colocalize *in vivo*. For example, our dataset’s clustering of the heavy and light ribosome subunits and the proteasome supports the notion that the fractionation preserves the colocalization of interaction networks. We also observe clusterization of different secretory compartments, especially the plasma membrane and endoplasmic reticulum (details about the clusters composition in control and treated samples are shown in the t-SNEs of **Figure S4**).

Altogether, these results indicate that the dataset is structured in a way that is dependent on subcellular localization, considering compartments delimited by membranes (i.e., organelles) and compartments delimited by protein complex formation (i.e., the nucleolus and the proteasome). This analysis provides a comprehensive investigation of HaCaT subcellular architecture, allowing for inferences about UVA-induced protein dynamic events. **UVA light elicits changes in the subcellular distribution of proteins**

Next, we used the TRANSPIRE pipeline (Kennedy et al., 2020) to classify possible UVA-triggered protein dynamic events in the spatial proteomics dataset. This pipeline has been developed for identification of protein translocations. TRANSPIRE creates synthetic translocation classes from organellar markers, trains a Gaussian process classifier based on the synthetic translocation classes and classifies translocations in the actual dataset. The basis of this approach relies on first concatenating the organellar markers between the different biological conditions to produce synthetic markers. Then the synthetic markers are further clustered to provide different translocation and non-translocation classes, allowing the algorithm to classify the directionality of protein trafficking across subcellular niches.

The algorithm performs all possible combinations of organellar markers between conditions to generate synthetic translocations of different classes. In this sense, “Nucleus to Cytosol” and “Mitochondria to Mitochondria” would represent two different classes. Thus, the algorithm’s output consists of the translocation classes attributed to each protein and translocation scores, calculated as described by Kennedy et al., 2020. False-positive rates (FPR) were calculated based on the learning model, and a stringent 0.1% FPR threshold was applied to define a true translocation event in our dataset. We tested multiple FPR thresholds and observed that for 0.1% and 0.01% FPR, we identified about the same number of translocations (**Figure S5A**). The output of the algorithm can be found in **Supplementary Spreadsheet S5**.

The classifier achieved a high level of accuracy during training (**Figure 5SB-D**) and in the test data, reaching F1 scores above 0.95 in the test set (as shown in **Figure 3A**). The classifier identified 217 possible targets of translocation (FPR 0.1%) altogether. The number of proteins assigned to each translocation class by the algorithm is shown in **Figure 3B**, revealing a predominance of dynamic events involving mitochondrial proteins. Secretory organelles also seem to play an important contribution to the overall translocations, secondly to mitochondria. By aligning the translocation classes in a circular plot (**Figure 3C**), it is possible to see that they are not equally distributed across the four subcellular niches. Indeed the efflux is more intense for mitochondria, in the direction of secretory compartments. This observation possibly reflects the crucial role secretory organelles play in protein trafficking between different subcellular niches and the importance of mitochondria in the stress response to radiation. Furthermore, the predominance of translocations from mitochondria to secretory organelles seems to be consistent with the partial approximation between the mitochondrial and secretory clusters in the three-dimensional t-SNE subcellular map of irradiated cells compared to controls (**Figure 2C**). Translocating proteins are significantly enriched for biological processes related to cellular localization (“cellular localization”, “establishment of localization in cell’, “cellular component organization”, “intracellular transport”). (**Figure 3D**).

**Figure 3.**
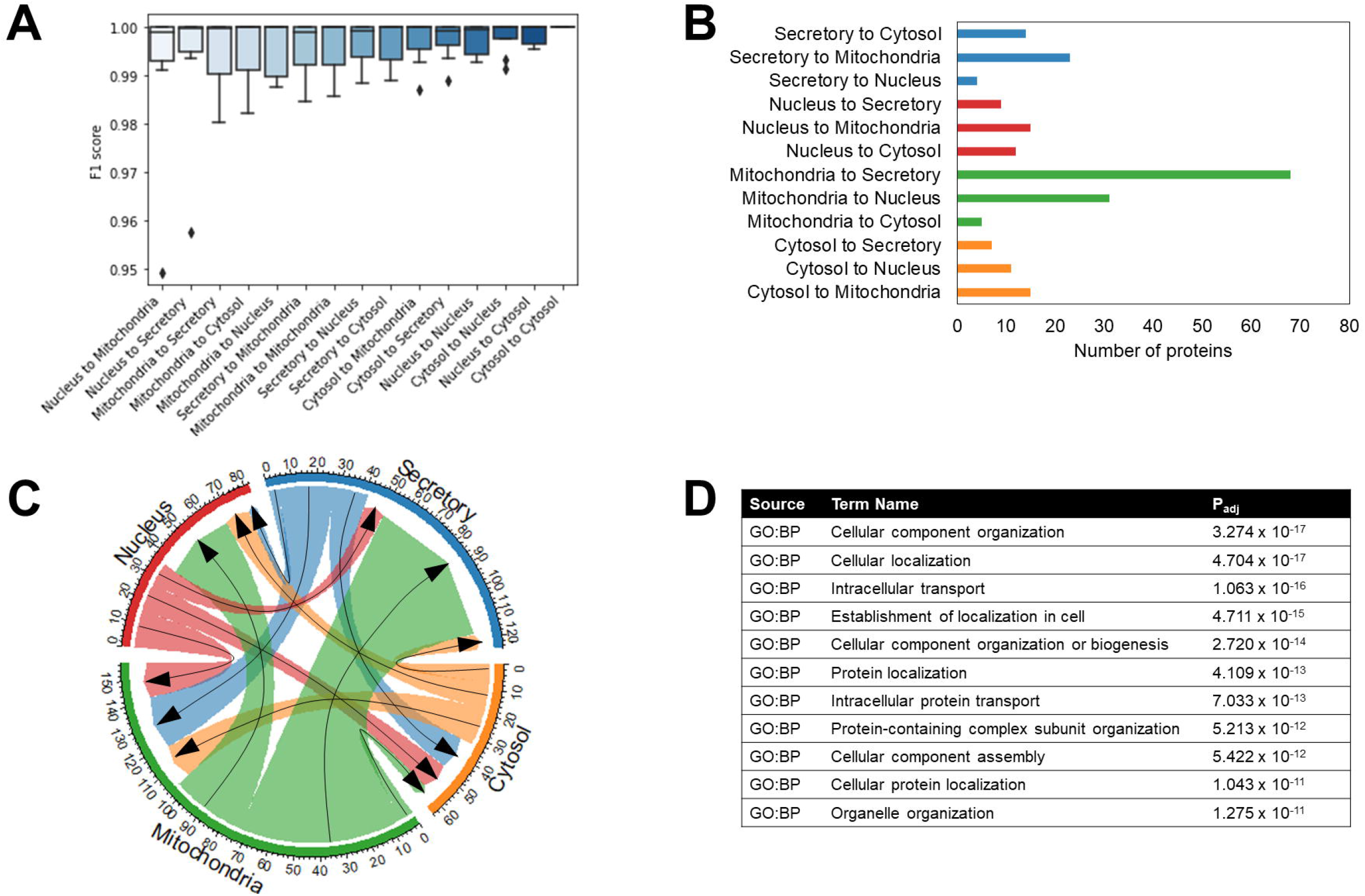
Prediction of UVA-induced translocations in HaCaT cells. **(A)** F1 scores obtained by the classifier in the test set. **(B)** Number of proteins assigned to each translocation class by the algorithm. **(C)** Circular plot representing UVA-induced translocations between subcellular niches, as identified by the classifier. **(D)** Enrichment analysis of the translocation targets based on Gene Ontology terms (BP: biological process).

Considering the list of possible translocation and the clusterization of organellar markers in the t-SNE plot, we performed a few validations by confocal microscopy. As DNA damage and oxidative stress are important and well-known triggers of UVA signaling in skin cells, we validated a nucleocytoplasmic translocation triggered by DNA damage to unequivocally confirm that DNA damage is an underlying feature of UVA-induced cellular stress response, even for a considerably low dose exposure. As mitochondrial proteins accounted for the majority of protein dynamic events, we also validated changes in mitochondrial dynamics associated to the shift of the mitochondrial cluster in the t-SNE plots between biological conditions. Overall, these validations emphasize DNA damage and metabolic stress as underlying features of the subcellular reorganization promoted by a low dose (6 J/cm^2^) of the least energetic component of the solar UV radiation in human keratinocytes.

### Spatial remodeling involves mitochondrial and DNA damage signaling

We first focused on curating the translocation labels classified by the TRANSPIRE algorithm using the neural networks classifications of subcellular localization in control and treated samples to validate a more stringent list of possible UVA-triggered translocations. Altogether, the translocations detected by using the neural networks algorithm were mostly covered by TRANSPIRE (**Figure 4A**). TRANSPIRE detects more targets than the neural networks algorithm (**Figure 4A**), which may be due to differences in the conception of the algorithms. For example, multilocalized proteins likely receive low confidence scores and are filtered out from the neural networks classifications, while TRANSPIRE is relatively agnostic in detecting translocations for multilocalized proteins (Kennedy et al., 2020). However, we observe that for the more strigent list of translocating proteins identified by both TRANSPIRE and the neural networks, we obtain the same pattern of migration between subcellular compartments as for the TRANSPIRE results as a whole (**Figure 4B**). The more stringent list of possible translocating proteins detected by both TRANSPIRE and the neural networks can be found in **Supplementary Spreadsheet S6**, together with enrichment analysis of this set of proteins. From this list, we chose one nucleocytoplasmic translocation well-known to be involved in the DNA damage response for immunofluorescence validation.

**Figure 4.**
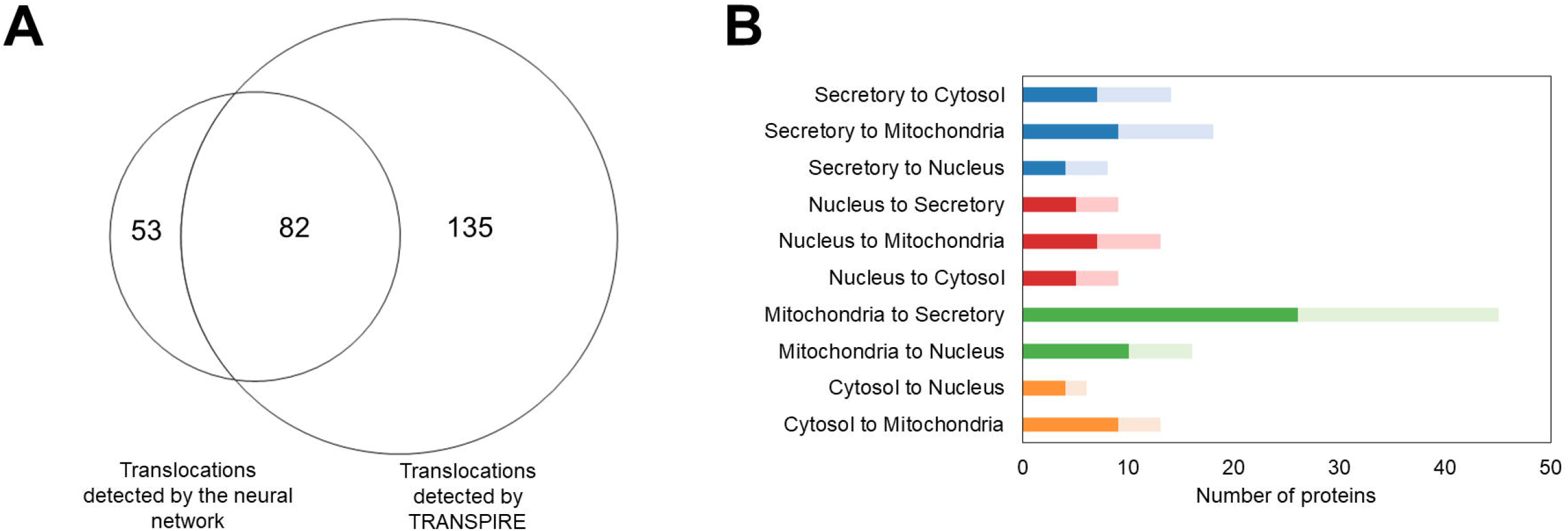
Comparison of the translocations classified by TRANSPIRE and neural networks algorithms. **(A)** Venn diagram representing the possible translocating proteins identified by each algorithm. **(B)** Bar plot representing the translocation classes classified by the algorithm. The darker shade of bars represents classifications obtained through the use of the neural networks and the lighter shade represents TRANSPIRE classifications.

The β subunit of CSNK2 (CSNK2B) was first implicated in the DNA damage response through its interaction with the tumor suppressor p53 (Filhol et al., 1992). CSNK2 is also involved in the phosphorylation of two NER components (XPB, CETN2) (Coin et al., 2004; Grecu and Assairi, 2014) and its translocation to the nucleus is associated to cell cycle arrest (Tripodi et al., 2007). Additionally, it has been demonstrated that XPC- and XPD-deficient cells expressing higher levels of CSNK2B are more resistant to UV-induced death (Teitz et al., 1990), especially since increases in CSNK2B lead to dramatic increases in CSNK2 activity (Cochet and Chambaz, 1983). In our experiment, CSNK2B shifts from a central position in the cytosolic cluster in controls to the interface between the cytosolic and nuclear clusters in irradiated samples (**Figure 5A**). This behavior is consistent with a significant difference between groups observed for this protein in the profile plot, especially in the last fraction, which is enriched with cytosolic proteins (**Figure 5B**). Redistribution of CSNK2B from the cytoplasm to the nucleus upon irradiation was corroborated by immunofluorescence, indicating that UVA exposure leads to the translocation of cytosolic CSNK2B to the nucleus (**Figure 4C**).

**Figure 5.**
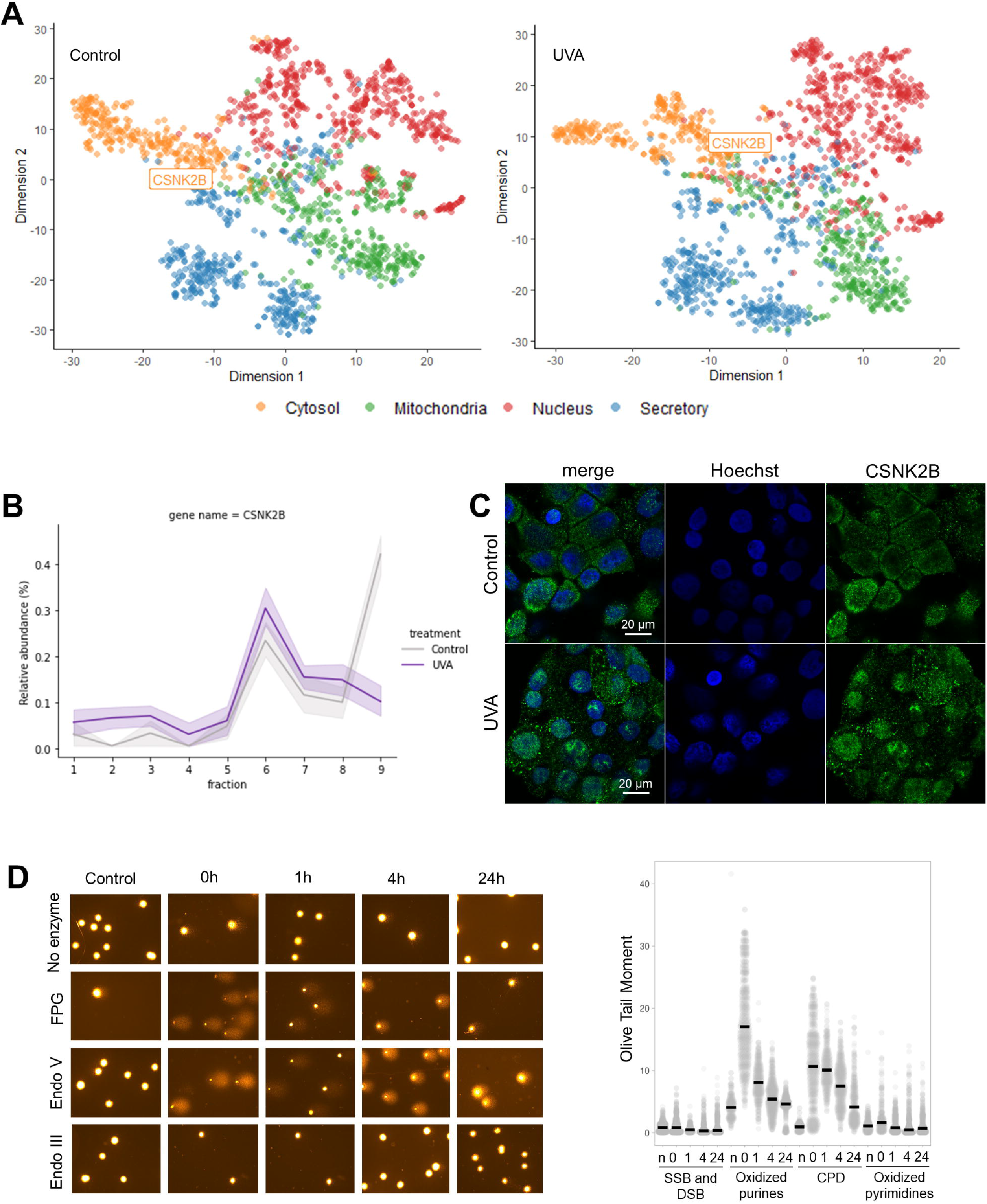
CSNK2B translocates to the nucleus upon DNA damage induced by UVA exposure. **(A)** 2D t-SNE plots representing the migration of CSNK2B from the cytosolic cluster in control samples to the nuclear cluster in UVA-irradiated cells. Colors represent the translocations predicted by the classifier (organelle of origin in controls and destination in UVA plot). **(B)** Profile plot obtained for CSKN2B in controls and irradiated samples. Lines represent the means of relative abundance, and shadowed intervals represent the standard errors. **(C)** Representative immunofluorescence images showing CSNK2B translocation from the cytosol to the nucleus after exposing HaCaT cells to UVA light. CSNK2B was immunostained (green), and the nucleus was stained with Hoechst (blue). Three independent replicates per group were analyzed. The bars indicate the 20 μm scale. **(D)** Comet assay results for control and irradiated cells. Representative images of randomly scored comets in slides from all conditions are represented on the left. The graph shows the semi-quantification of each type of DNA damage over time. Points represent Olive Tail Moments scored for all measured cells, and black bars represent the medians of all points (n = 4 independent experiments). In this case, CPD quantification represents the sum of CPDs and abasic sites. The number of abasic sites is at most equal to the oxidized purine number. Subtracting the averages of oxidized purines from the CPD quantification data still leaves a CPD repair curve with the same general time-course.

To confirm that our irradiation conditions generated significant levels of DNA damage, we performed a modified version of the comet assay to detect different types of DNA lesions in cells following exposure to 6 J/cm^2^ of UVA light (**Figure 4D**). The comet assay was modified through the addition of formamidopyrimidine-DNA glycosylase (FPG), endonuclease V (endoV) and endonuclease III (endoIII) to detect oxidized pyrimidines, CPDs and oxidized purines, respectively. The predominant types of lesions generated immediately after exposure to UVA are CPDs and oxidized purines, in agreement with what has been previously described for this radiation dose (Delinasios et al., 2018). However, while oxidized purines seem to be efficiently removed from the DNA one hour after exposure to the radiation. CPDs begin to be repaired soon after irradiation and require more than 24 hours to be completely repaired. The DNA lesion profile identified here and its repair kinetics are consistent with NER activation, and thus consistent with triggering of translocation events associated to this pathway. Even though UVA generates lower levels of CPD than UVB, CPD generation can still promote CSNK2B recruitment to the nucleus, leading possibly to cell cycle arrest and allowing for DNA repair.

### UVA light promotes metabolic stress through mitochondrial fragmentation

Changes in organelle dynamics or morphology may yield hypothesis on their functional states, while the distribution of proteins over subcellular niches hints to organelle or pathway activity (Palla et al., 2022). In this sense, cellular phenotypes can be influenced both by organelle properties (i.e., morphology and dynamics) and by the distibution of molecules over the cell. Supervised learning algorithms applied to spatial proteomics datasets are designed to detect translocations of proteins between subcellular niches (Gatto et al., 2014b; Kennedy et al., 2020). Conversely, changes in organelle dynamics are more likely to lead to systematic alterations in the fractionation profiles of all proteins localized in the affected organelle. This happens because changes in organelle morphology and dynamics may lead to differential sedimentation patterns of the whole organelle during differential centrifugation.

The first evidence we found of a change in mitochondrial dynamics involves the shift of the mitochondrial cluster in the subcellular t-SNE map of irradiated samples relative to controls (as previously shown in **Figure 2B** and **Figure S1E-F**). Following this finding, we investigated possible alterations in the fractionation profiles of mitochondrial markers, as an evidence of broader changes in the fractionation profiles of mitochondrial proteins, which would not be detected by TRANSPIRE or the neural networks algorithms. For this purpose, the fractionation profiles of organellar markers in control samples were overlaid with the fractionation profiles of organellar markers in UVA-irradiated samples (**Figure 6A**). These plots reveal that the fractionation profile of mitochondrial markers in irradiated samples display a subtle shift relative to controls. Other comparments do not show change in fractionation profiles comparing UVA and non-irradiated samples. Even though the shift in the fractionation profile of mitochondrial markers is subtle, it is consistently observed across many structural mitochondrial proteins (as exemplified in **Figure 6B**).

**Figure 6.**
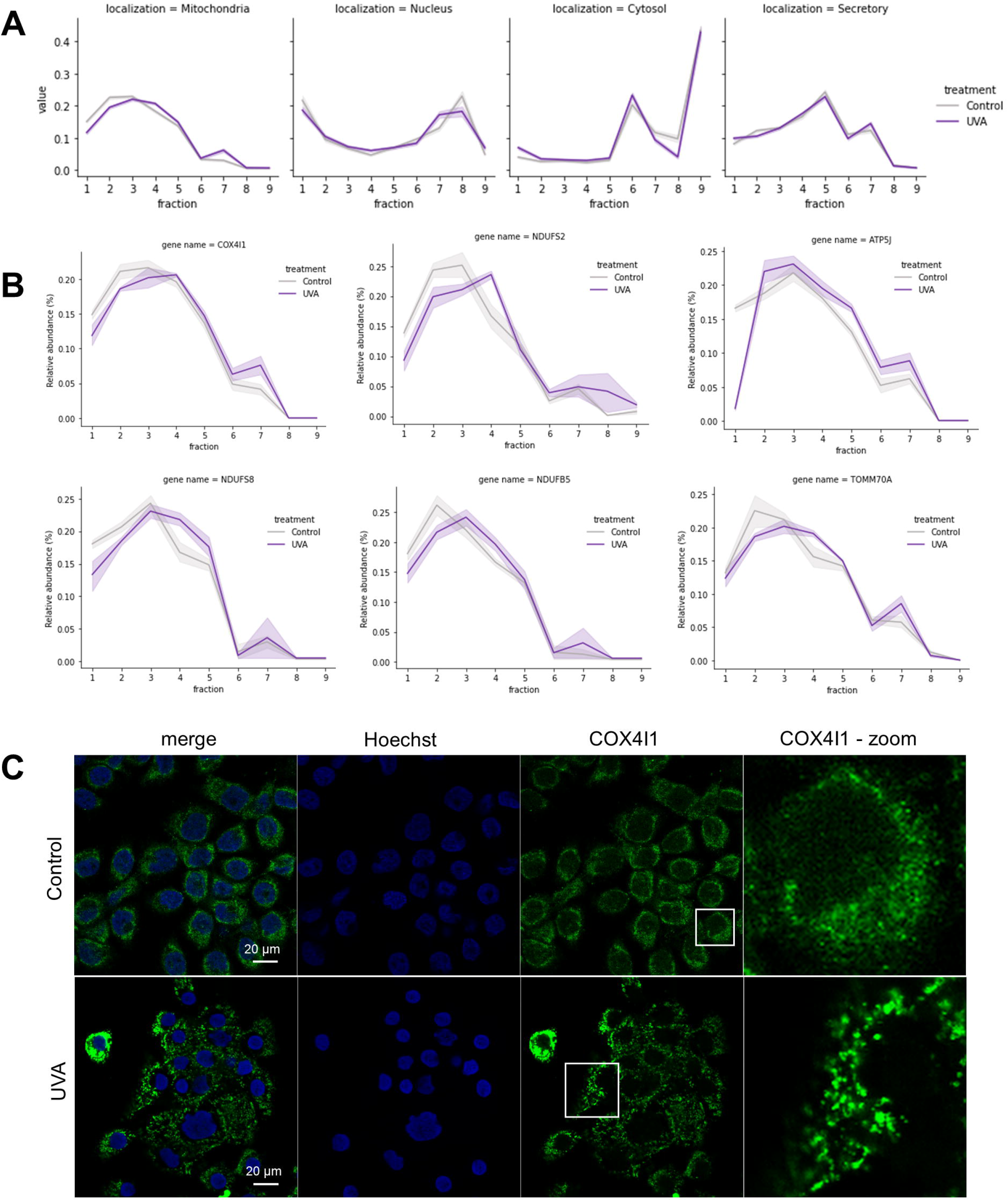
UVA induces changes in mitochondrial morphology that are associated with changes in protein fractionation. **(A)** Profile plots of organellar markers in control and irradiated samples. Lines represent the means of relative abundance, and shadowed intervals represent standard errors. **(B)** Profile plots of representative proteins from the mitochondrial membrane. Lines represent the means of relative abundance, and shadowed intervals represent standard errors. **(C)** Immunostaining of COX4I1 (green). The nucleus was stained with Hoechst (blue). Two biologically independent experiments were performed, with similar results obtained. The bars indicate the 20 μm scale.

To confirm that the alterations in the fractionation profiles of mitochondrial markers between conditions might represent alterations in mitochondrial morphology and dynamics, we immunolabeled a respiratory chain component (COX4I1), which is not expected to translocate, and performed an IF experiment. As shown in **Figure 6C**, in controls, COX4I1 displays the typical tubular, interconnected appearance of the mitochondrial network, which can be observed through the diffuse aspect of the fluorescence labeling, and forms punctate structures in irradiated samples, a sign of UVA-induced mitochondrial fragmentation (as has been similarly observed by Kowaltowski et al., 2019).

To determine if changes in the fractionation profiling of non-translocating and non-structural mitochondrial proteins could also reflect mitochondrial fragmentation, we investigated the spatial redistribution of fumarase (FH) and ornithine aminotransferase (OAT) in irradiated cells. OAT and FH were not detected as translocations, considering the stringent 0.1% FPR threshold we used in the TRANSPIRE analysis. We also monitored PDHA1 in the same experiment to check for colocalization of structural (PDHA1) and non-structural mitochondrial (OAT and FH) proteins, since PDHA1 is a structural mitochondrial protein and also not expected to translocate. Importantly, FH and OAT display similar fractionation patterns (**Figure 7A**). Both proteins display decreasing levels in the first fractions (1-3) of irradiated samples compared to controls, accompanied by increased levels in the last fraction (**Figure 7A**). Immunofluorescence images show that OAT and FH display the same pattern of subcellular localization as PDHA1, indicating that translocation to other subcellular compartments does not seem to be occuring. Instead the fluorescence pattern confirmed the same mitochondrial fragmentation phenomenon observed for labeling of structural mitochondrial proteins (COX4I1 and PDHA1), reinforcing our previous results (**Figure 7B-C**). These results demonstrate that, even though the learning algorithms detected translocations of mitochondrial components, there is another layer of biological information in our spatial proteomics dataset: systematic changes in the fractionation profiles of mitochondrial markers are a consequence of mitochondrial fragmentation.

**Figure 7.**
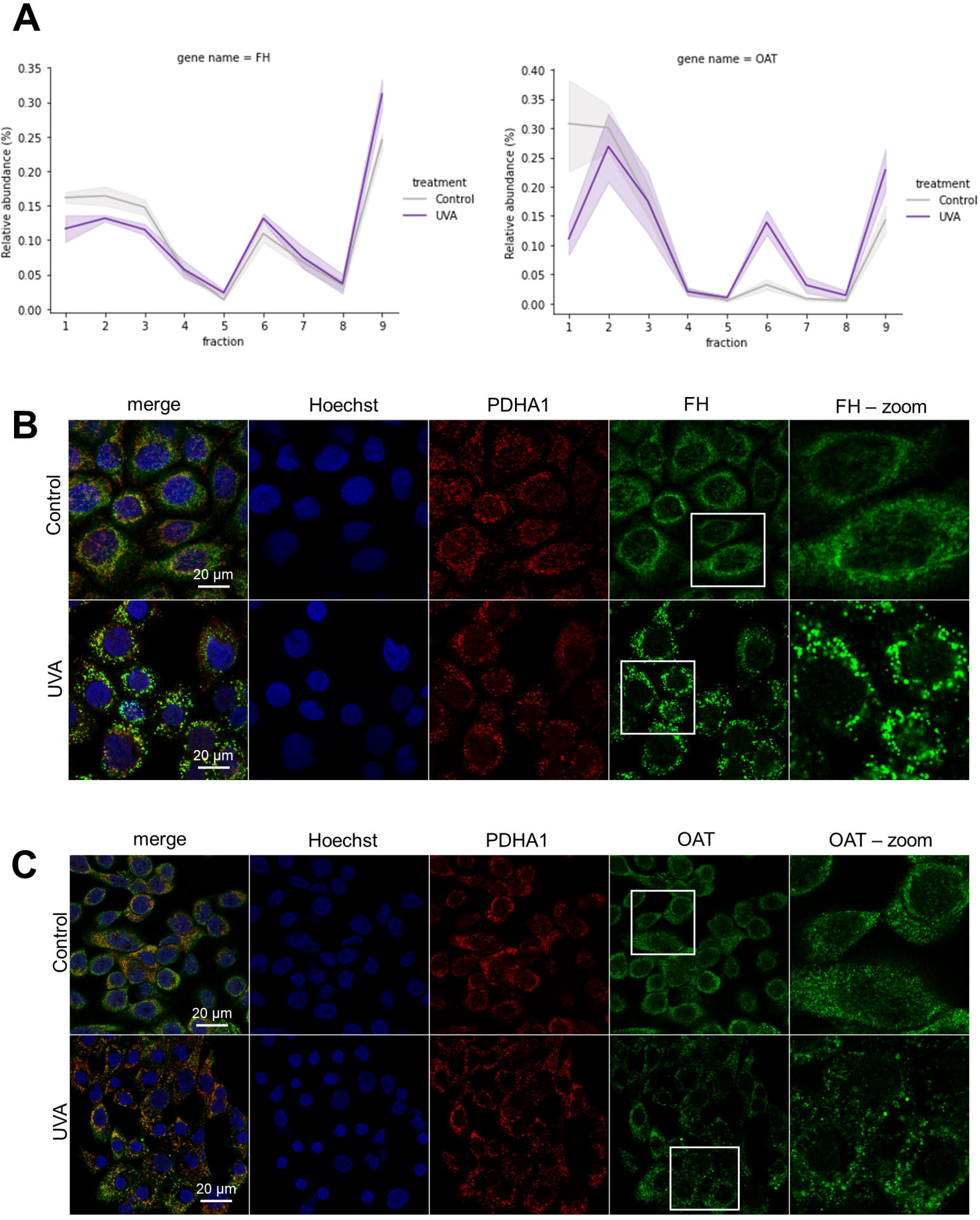
Changes in the fractionation profiling of proteins from the mitochondrial matrix in UVA-exposed cells compared to controls reflect mitochondrial fragmentation. **(A)** Profile plots obtained for FH and OAT in control and irradiated samples. Lines represent the means of relative abundance, and shadowed intervals represent standard errors. **(B)** Immunostaining of FH (green) in HaCaT cells exposed to UVA or mock-treated. PDHA1 (red) was immunolabeled as a structural mitochondrial marker. The nucleus was stained with Hoechst (blue). Three independent experiments were performed, and similar results were obtained. **(C)** Immunostaining of OAT (green) in HaCaT cells exposed to UVA or mock-treated. Similarly, PDHA1 (red) was used as a mitochondrial marker, and the nucleus was stained with Hoechst (blue). Three independent experiments were performed, and similar results were obtained. The bars indicate the 20 μm scale.

Since cells displaying fragmented mitochondria usually have a reduced respiratory capacity (Sabouny and Shutt, 2020), we measured oxygen consumption rates in HaCaT cells exposed to UVA light using a Seahorse Analyzer XF24 to validate the functional impact of mitochondrial fragmentation. Accordingly, basal and maximal mitochondrial respiration are decreased in irradiated cells compared to control samples, supporting the notion of electron transport chain dysfunction (**Figure 8A**). Changes in mitochondrial respiration were accompanied by a decrease in the cell’s reductive power up to 24 hours after irradiation (MTT results, **Figure S6**), without losses in viability, as inferred by the trypan blue exclusion assay (**Figure S6**). However, we are careful to point out that we did not perform an specific assay for apoptotic cells. The reduction in the cell’s reductive power occurs in a radiation dose-dependent manner (**Figure S6**)..

**Figure 8.**
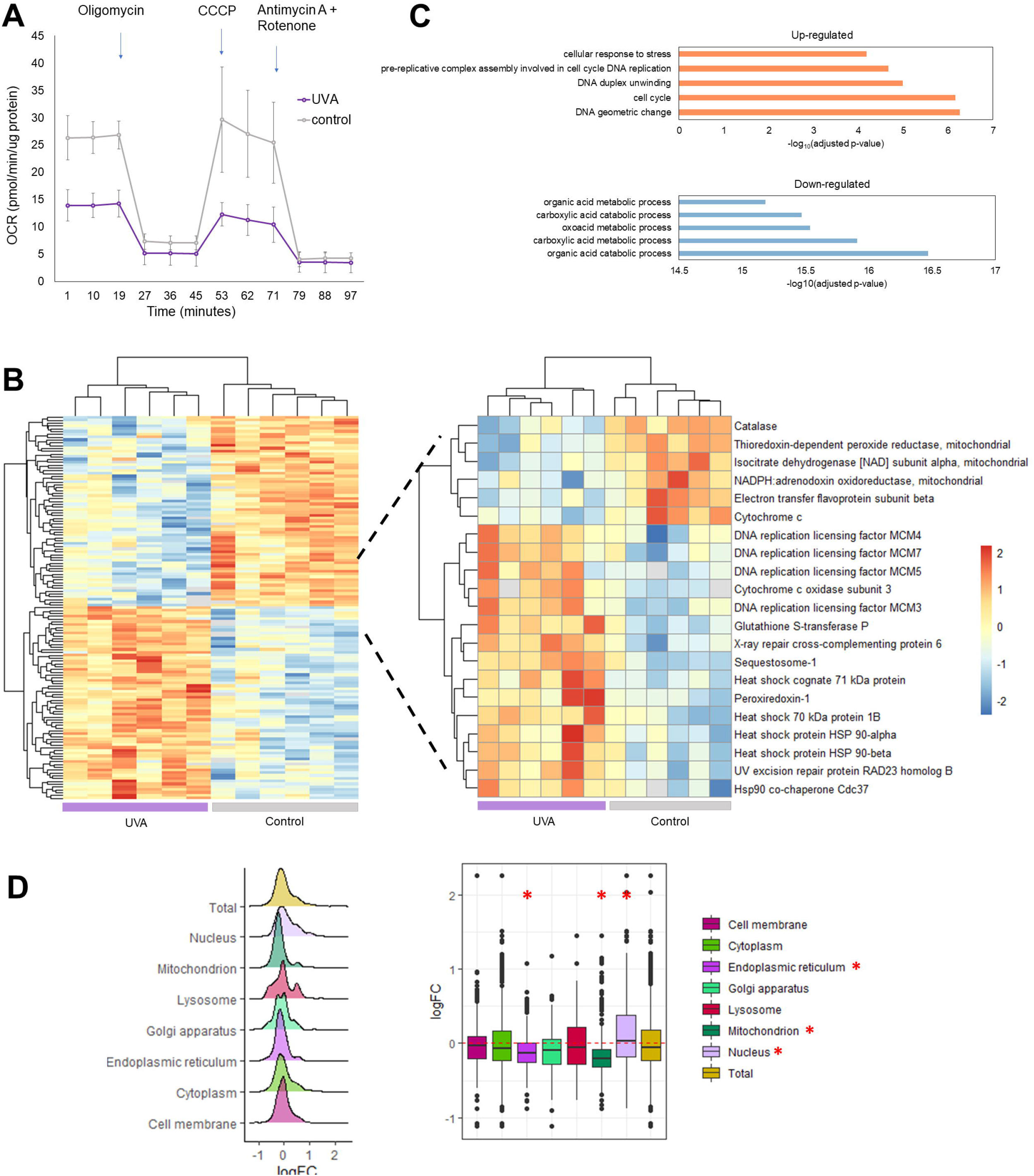
UVA-induced changes in mitochondrial dynamics impact respiratory function. **(A)** Oxygen consumption rates (OCR) were measured in irradiated and control cells before and after the addition of 1 μM oligomycin, 1 μM CCCP and a solution containing 1 μM antimycin and 1 μM rotenone (n = 4 biological replicates). Points represent means and error bars represent standard deviations of biological replicates. **(B)** Hierarchical clustering of differentially regulated proteins comparing HaCaT cells exposed to UVA versus controls (Student’s T-test, 0.05 FDR correction). The color gradient represents z-scored LFQ intensities, and columns represent replicates (n = 6 per group). Stress-responsive proteins are highlighted. **(C)** Enrichment analysis of differentially abundant proteins according to GO terms. **(D)** Compartment-specific proteome changes in irradiated versus control HaCaT cells one and a half hours after UVA exposure (n = 6). The analysis of the log_2_ fold changes of irradiated HaCaT cells in relation to controls was performed according to (Parca et al., 2018): proteins were assigned to compartments according to GO-terms and each compartment was tested for significant difference against the whole proteome (Wilcoxon rank sum test with 0.5% FDR correction). The asterisks indicate compartments with corrected p-values lower than 0.05.

Since UVA light is known to cause oxidative and genotoxic stresses (Schuch et al., 2017), we tested if a 6 J/cm^2^ dose of UVA could promote alterations in the levels of stress-responsive proteins one and a half hour after radiation exposure. We opted for performing the total proteome analysis at a later time point than the spatial proteomics experiment because we reasoned that alterations in protein levels due to synthesis/degradation would be a longer-term response compared to early signaling dynamic events. We also chose this time point because we intended to look at changes in protein levels that resulted from the mitochondrial and DNA damage we detected previously via the comet assay and respirometry. After this period, 138 proteins were significantly modulated between groups (**Figure 8B**), the general enrichment analysis of up and down-regulated proteins is shown in **Figure 8C**. Focusing on stress-responsive proteins (**Figure 8B**), we observed the up-regulation of DNA damage response components (RAD23B and XRCC6), a few DNA replication licensing factors, antioxidant enzymes (GSTP1 and PRDX1) and heat shock proteins. Additionally, a few subunits of the electron transport chain complexes and a few redox-responsive proteins (CAT and PRDX3) were down-regulated, possibly due to depletion, as has been already described for cell under stress induced by UVA radiation (Maresca et al., 2006). This dataset and the enrichment analysis of differentially abundant proteins are provided in **Supplementary Spreadsheet S7**. Additionally, the lists of proteins exclusively present in one biological condition are provided in **Supplementary Spreadsheet S8**.

By analyzing the fold change of proteins between treatments in a compartment-specific fashion (**Figure 8D**), we found that the fold change of mitochondrial proteins is significantly lower when compared to the whole proteome (p = 1.47 × 10^-17^, Wilcox test, FDR correction), suggesting that decreasing levels of electron transport chain components recapitulate mitochondrial proteome changes as a whole. Importantly, mitochondrial fragmentation usually facilitates mitophagy of damaged mitochondria (Twig and Shirihai, 2011).

These results show that exposing skin cells to UVA light impacts mitochondrial dynamics, leading to fragmentation, respiratory dysfunction, and the upregulation of stress response proteins.

## Discussion

The present study is the first to provide a map of subcellular protein reorganization induced by the UVA component of sunlight in a skin cell type. High sensitivity MS-based proteomics coupled to machine learning algorithms quantified and assigned subcellular localization for over 1600 proteins in human keratinocytes exposed to UVA light. Our unbiased approach revealed that a single low dose of UVA light could affect the proteomic architecture of skin cells, provoking the reorganization of subcellular structures due to genotoxic and metabolic stresses.

In this work, about 12% of the identified and quantified proteins (over 200 proteins from a total of 1600) relocalized in response to UVA exposure. Our results showed that redistribution of proteins across subcellular niches encompass different phenomena, such as changes in organelle dynamics and translocations. After considering all redistribution events, important modulators of cellular metabolism, mitochondrial function, protein trafficking, signaling pathways and DNA damage response were identified.

Previously it was reported that DNA damage response rewires metabolic circuits, fine-tuning protein synthesis, trafficking and secretion (Chatzidoukaki et al., 2020). However, it is not clear how genotoxic components of the sunlight affect protein localization or organelle architecture and interactions. In this work we confirmed that UVA exposure caused a nucleocytoplasmic dynamic event induced by DNA damage. For instance, our algorithms detected with high confidence the nucleocytoplasmic translocation of CSNK2B in UVA-irradiated cells, a finding further confirmed by confocal microscopy. CSNK2 has many biological targets, maintaining cellular viability and participating in the DNA damage response (Gray et al., 2014; Montenarh, 2016; Yefi et al., 2011). Its role in the cellular response against UVR has been described in terms of its interaction with p53 and NER components as well as by its role in cell cycle arrest (Montenarh, 2016). Indeed, using the comet assay, we observed that UVA radiation leads to simultaneous CPD formation and CSNK2B translocation, as expected. We also monitored DNA damage over time and observed that CPDs are repaired over 24 hours, indicating NER activation. Collectively, these results demonstrate that UVA triggers a classical DNA damage signaling pathway, even though it generates lower levels of CPD than the more energetic UVB light.

The most striking result of our systematic proteomic profiling was identifying mitochondria as one of the main targets of UVA-induced stress. In our experiment, we detected two types of dynamic events: redistributions of mitochondrial proteins over the subcellular space, which were detected by the TRANSPIRE and neural networks algorithms, as well as changes in mitochondrial dynamics, which were reflected as systematic changes in the fractionation profiles of mitochondrial markers. We showed that UVA induces mitochondrial fragmentation, up-regulates redox-responsive proteins and reduces the respiratory rate, leading to changes in the cells’ overall energetic status. Mitochondrial fragmentation depends on the balance between fusion and fission, enabling adaptation to stress and different metabolic demands (Sprenger and Langer, 2019). If fragmentation of the interconnected mitochondrial network occurs transiently, it can facilitate mitophagy, for example, leading to adaptation to stress (Sprenger and Langer, 2019). For instance, mitochondrial fragmentation and mitophagy are essential for keratinocyte cornification, a physiological process (Simpson et al., 2021). On the other hand, chronic fragmentation may be associated to other types of cell death and tissue damage (Sprenger and Langer, 2019). Thus, for example, it has been demonstrated that healthy and cancerous skin display different patterns of mitochondrial organization and morphology (Pouli et al., 2016).

These results expand on previous characterizations of mitochondrial dysfunction in response to UV radiation (Djavaheri-Mergny et al., 2001; Jugé et al., 2016) and show that alterations occur even with acute low-dose exposures to UVA, the least energetic component of the UV spectrum. It has been suggested that UVA-induced deletions in mtDNA underlie the long-term effects of UVA during photoaging (Berneburg et al., 2004). However, our results suggest that UVA also has short-term effects on the mitochondria, acting as a potent stressor immediately after exposure. Some endogenous metabolites have been proposed to play a role in UVA’s photosensitization in skin cells, such as flavin-derivatives, NADH, NADPH, FADH, urocanic acid, porphyrins and some sterols (Wondrak et al., 2006). Mitochondria, in particular, contain high concentrations of putative UVA chromophores, such as flavin-derivatives, NADH, FADH and NADPH, which could mediate the damage to this organelle.

Several studies showed that high doses of UVB irradiation (e.g., > 100 mJ/cm^2^) trigger mitochondrial fragmentation in keratinocytes (Jugé et al., 2016; Wang et al., 2015, p. 1; Zhang et al., 2016, p. 1). On the other hand, UVC (60 mJ/cm^2^) leads to mitochondrial hyperfusion instead of fragmentation in mouse fibroblasts, suggesting that UVR-induced modulations of mitochondrial dynamics are complex and context-dependent (Tondera et al., 2009). Our results show that even a low dose of the less energetic UVA light is capable of inducing mitochondrial fragmentation. UVB-induced mitochondrial fragmentation is dependent on DRP1 mitochondrial translocation, with partial roles for MFN1 and OPA1 (Jugé et al., 2016), frequently followed by apoptosis (Wang et al., 2015; Zhang et al., 2016). In our experiments with UVA irradiation, we did not detect strong evidences of attenuated cell viability, as inferred by the trypan blue assay.

In summary, our dataset provides valuable information about UVA-triggered translocation events in subcellular niches. Our experimental strategy employing cellular fractionation, MS-based proteomics and machine learning algorithms revealed UVA redistributed approximately 12% of the skin cell proteome, highlighted by the up-regulation of redox-responsive proteins, DNA damage and mitochondrial fragmentation. Overall, our dataset opens up possibilities for further investigation of UVA-triggered protein dynamic events in less studied subcellular niches

### Limitations of the study

Our work has some limitations. First, this study is based on a shotgun, label-free proteomics quantification approach. Data-dependent acquisition is well-known to generate a high number of missing values in proteomics datasets, due to the semi-stochasticity of ion fragmentation. Here we dealt with the limitations of this technique by filtering part of the missing values and by using an imputation method that takes into account both values missing at random and not at random. Using isotopic or isobaric labelling, for example, could lead to acquisition of a more complete set of protein quantifications and consequently to more accurate classifications of subcellular localization and translocations. Furthermore, another limitation of our study includes the limited number of target validations. Microscopy validation is a valuable tool to confirm and extract mechanistic information of the possible translocation targets. More extensive, high-throughput microscopy validations of the possible translocation targets classified by the machine learning algorithms can reveal new players in early skin tumorigenesis and early stress signaling induced by UVA light.

## Acknowledgments

We thank professor Marisa Helena Gennari de Medeiros for kindly providing access to her laboratory and equipment. We are grateful to professor Fabio Rodrigues and members of his laboratory, especially Evandro Pereira da Silva, for kindly helping with the solar simulator. We also thank Camille C. Caldeira da Silva and professor Alicia Kowaltowski for kindly helping with the Seahorse experiment. We are grateful to Dr. Mariana Pereira Massafera and MSc. Izaura Nobuko Toma for their technical assistance. The Redox Proteomics Core of the Mass Spectrometry Resource at Chemistry Institute, University of Sao Paulo, is acknowledged for access to state-of-the-art MS instrumentation. The Albert Einstein Hospital is also acknowledged for technical assistance and access to the confocal microscope. We thank Fundação de Amparo à Pesquisa do Estado de São Paulo (FAPESP) (grants 2012/12663-1, 2016/11430-4, 2016/00696-3, 15/07768-7), CEPID Redoxoma (2013/07937-8), Conselho Nacional de Desenvolvimento Científico e Tecnológico (CNPq) (grants 402683/2016-1 and 302120/2018-1), Coordenação de Aperfeiçoamento de Pessoal de Nível Superior (CAPES), NAP Redoxoma (PRPUSP—2011.1.9352.1.8) and John Simon Guggenheim Memorial Foundation (PDM Fellowship) for financial support. HPV is recipient of a FAPESP fellowship (2016/11430-4).

## Authors Contributions

HPV, GER and PDM conceptualized the study. HPV performed experiments, analyzed the data and wrote the first draft of the manuscript. FGR and APYC performed the immunofluorescence assays and discussed the results. BDCC performed the comet assay experiment and also discussed the results. GER and PDM acquired funding, provided the resources, critically read and edited the final version of the manuscript.

## STAR METHODS

### RESOURCE AVAILABILITY

#### Lead contact

Further information and requests for resources and reagents should be directed to and will be fulfilled by the lead contact: Prof. Dr. Paolo Di Mascio.

#### Materials availability

This study did not generate new materials or new unique reagents.

#### Data and code availability

- All data reported in this paper will be shared by the lead contact upon request. The raw MS files are publicly available as of the data of the publication in the PRIDE repository. The identifier is listed in the key resources table.
- This paper does not report original code.
- Additional information required to reanalyze the data reported in this paper is available from the lead contact upon request.

### EXPERIMENTAL MODEL

#### Keratinocyte cell line

HaCaT cell line (caucasian, male), a spontaneously immortalized human keratinocyte, was cultured in 5% CO_2_ at 37°C and grown in Dulbecco’s Modified Eagle’s Medium (DMEM) supplemented with 10% fetal bovine serum, 100 U/mL penicillin and 100 μg/mL streptomycin. Professor Mauricio S. Baptista (Institute of Chemistry, University of São Paulo) provided the cell line, and it was tested for mycoplasma contamination. The cell line was not authenticated.

HaCaT cells are commonly used as a pre-malignant model (Fusenig and Boukamp, 1998). HaCaT cells do not harbor any viral sequences (Boukamp et al., 1997; Mueller et al., 2001) and were probably immortalized as a consequence of UV-like mutations in p53 (Lehman et al., 1993). This cell line presents other characteristics of an initiated cell line, such as increased telomerase activity, silencing of p16, and defective regulation of p21 expression and function (Ren et al., 2006). Despite these alterations, HaCaT cells have a stable chromosome content and remain non-tumorigenic (Boukamp et al., 1997). As our experiment required a large number of cells, we opted for using an immortalized cell line, but the genetic status of this cell lineage is well-defined. Cells were used between passages 13 and 17.

### METHOD DETAILS

#### Irradiation conditions

An Oriel SOL-UV 2 solar simulator (Newport, USA) equipped with a Xenon arc lamp was used for cell irradiation. The simulator was equipped with an IR bandpass blocking filter (UG11 Glass, model FSQ-UG11, Newport, MA, USA) plus a UVB-blocking filter (320 nm Cut-on, model SOL-UV-A-F, Newport, MA, USA). Emission spectra of the simulator radiation with and without the UVB-blocking filter are displayed in **Figure S7**. Before irradiation, the simulator’s output was measured with a dosimeter from International Light Inc (Newburyport, MA, USA), model IL1700, with a SED033 detector. Using the IR and UVB blocking filters, the output measured in the area where the cell plates would be positioned, at a 10 cm distance from the light source, yielded a mean of 5.0 mW/cm^2^, with a maximum variation of 10% between biological replicates. Each dish was irradiated for 26 minutes, corresponding to a total dose of 6 J/cm^2^, which humans can be exposed to during routine daily living without affecting cellular viability (**Figure S6**). All the assays described in this study were performed under the same UVA dosage (6 J/cm^2^). Cells were washed three times with phosphate-buffered saline (PBS) and kept in PBS during irradiation (26 minutes). Controls were kept in PBS and maintained in the dark at room temperature for the same amount of time.

#### Subcellular proteome sample preparation

For the spatial proteomics assay, two million cells were plated in 100 mm dishes 48 hours before the experiments (until cells reached 80-90% confluency). An entire dish containing around eight million cells yielded at least 10 μg of protein in the fraction with the lowest yield, which was enough for mass spectrometry analysis.

Cells were trypsinized and harvested by centrifugation 30 minutes after irradiation. The cell pellet was washed twice in PBS and incubated for 10 minutes in 1 mL of hypotonic lysis buffer (25□mM Tris-HCl, pH 7.5, 50□mM Sucrose, 0.5□mM MgCl_2_, 0.2□mM EGTA) on ice. Cells were then transferred to a Potter-Elvehjem homogenizer and homogenized with 30 strokes on ice (until at most 70% of cells were stained with trypan blue). After homogenization, 110 μL of hypertonic sucrose buffer (2.5□M sucrose, 25□mM Tris pH 7.5, 0.5□mM MgCl2, 0.2□mM EGTA) was used to restore osmolarity. The cell lysate was transferred to 2 mL tubes and centrifuged twice at 200 × *g* for 5 minutes to remove intact cells. The lysate was then subjected to a series of differential centrifugations: 1000 × *g* for 10 minutes, 3000 × *g* for 10 minutes, 5000 × *g* for 10 minutes, 9000 × *g* for 15 minutes, 12000 × *g* for 15 minutes, 15000 × *g* for 15 minutes, 30000 × *g* for 20 minutes and 80000 × *g* for 40 minutes. In total, each of the five biological replicates of each condition yielded nine fractions. The supernatant was collected because it contains the remaining cytosolic proteins. Afterward, fractions enriched with different organelles were lysed in 8 M urea containing 0.1% deoxycholate. The total protein concentrations were quantified using a BCA assay kit (Thermo Scientific), and 10 μg of protein per fraction were digested and analyzed by mass spectrometry.

#### Whole-cell proteome sample preparation

For whole-cell proteome analysis, 400,000 cells were plated in 6-well plates 24 hours before the experiments and incubation was followed by irradiation. One and a half hour after irradiation with 6 J/cm^2^ of UVA light, cells were washed five times with PBS, and scraped in 500 μL of a solution containing 100 mM ammonium bicarbonate, 8 M urea and protease inhibitors (cOmplete™ Protease Inhibitor Cocktail, Roche). Cell lysates were kept on ice for one hour and after that, they were precipitated overnight with 3 volumes of cold (−20 °C) acetone. Precipitated proteins were collected by centrifugation and acetone was discarded. Pellets were air-dried for about 10 minutes and resuspended in 100 mM ammonium bicarbonate buffer containing 8 M urea. Protein concentration was measured using a Pierce™ BCA Protein Assay Kit (Thermo Fisher Scientific) and 10 μg of protein per sample were digested and analyzed by mass spectrometry.

#### Protein digestion

Aliquots corresponding to 10 μg of protein per sample were reduced with 5 mM dithiothreitol for one hour, alkylated with 15 mM iodoacetamide for 30 minutes, diluted ten-fold with 100 mM ammonium bicarbonate, and digested by the addition of two aliquots of trypsin (1:40 and 1:50, respectively, with an interval of four hours between the additions). The samples were digested overnight at 30°C with agitation (400 rpm). Digestion was stopped by adding 4% trifluoracetic acid (TFA), and then the samples were dried. Samples were desalted using the StageTip protocol (Rappsilber et al., 2007). Peptides were washed ten times with 0.1% TFA in the StageTips and eluted with 50% acetonitrile and 0.1% TFA.

#### LC-MS/MS measurements

Each sample was injected in an Orbitrap Fusion Lumos mass spectrometer (Thermo Fisher Scientific, Bremen, Germany) coupled to a Nano EASY-nLC 1200 (Thermo Fisher Scientific, Bremen, Germany). Peptides were injected into a trap column (nanoViper C18, 3 μm, 75 μm × 2 cm, Thermo Scientific) with 12 μL of solvent A (0.1% formic acid) at 980 bar. After this period, the trapped peptides were eluted onto a C18 column (nanoViper C18, 2 μm, 75 μm × 15 cm, Thermo Scientific) at a flow rate of 300 nL/min and subsequently separated with a 5-28% acetonitrile gradient with 0.1% formic acid for 80 minutes, followed by a 28-40% acetonitrile gradient with 0.1% formic acid for 10 minutes.

The eluted peptides were detected in the data-dependent acquisition mode under positive electrospray ionization conditions. A full scan (*m/z* 400-1600) was acquired at a 60000 resolution, followed by HCD fragmentation of the most intense ions, considering an intensity threshold of 5 × 10^4^. Ions were filtered for fragmentation by the quadrupole with a transmission window of 1.2 *m/z*. HCD fragmentation was performed with a normalized collision energy of 30, and the Orbitrap detector analyzed the fragments with a 30000 resolution. The number of MS2 events between full scans was determined by a cycle time of 3 seconds. A total of 5 × 10^5^ and 5 × 10^4^ ions were injected in the Orbitrap with accumulation times of 50 and 54 seconds for the full scan and MS2 acquisition, respectively. Monocharged ions or ions with undetermined charges were not selected for fragmentation.

#### Comet assay

A total of 500,000 cells were plated in 6-well plates 24 hours before the experiment (n = 4). After irradiation, cells were trypsinized and collected by centrifugation. The supernatant was discarded, and cell pellets were mixed with 100 μL of PBS. 10 μL of cell suspension was added to 90 μL of 0.5% low melting point agarose. Subsequently, 75 μL of this cell suspension was pipetted onto slides pre-coated with 1.5% normal melting point agarose. Slides were covered with coverslips and kept at 4°C for 30 minutes to allow the agarose to solidify. Next, the coverslips were removed, and the slides were kept in a tank containing lysis buffer (2.5 M NaCl, 100 mM EDTA, 10 mM Tris, 1% Triton X-100, and 10% DMSO, pH 10) overnight at 4°C in the dark.

After lysis, slides were washed with cold PBS three times in the dark and immersed three times in cold reaction buffer (40 mM HEPES, 0.1 M KCl, 0.5 mM EDTA, 0.2 mg/mL BSA, pH 8) for 5 minutes each time. After that, the reaction buffer or reaction buffer containing T4 endonuclease V (0.1 U/mL), FPG (0.2 U/mL) or endonuclease III (10 U/mL) enzymes were pipetted onto each slide. Coverslips were placed over the slides, and they were incubated for 30 minutes at 37°C in the dark. The slides were then transferred to cold electrophoresis buffer (10 M NaOH, 200 mM EDTA, pH 13 in water) and incubated for 20 minutes. Then, the slides were submitted to electrophoresis for 20 minutes at 25 V, 300 mA. After electrophoresis, the slides were immersed in neutralizing solution (0.4 Tris, pH 7.5) three times for 5 minutes each time and fixed in methanol for 10 minutes. After washing, all the slides were air-dried at room temperature. The DNA was stained with 20 μL of a solution containing 2.5 ug/mL of propidium iodide for 10 minutes. Fifty randomly selected comets per sample were analyzed on a fluorescence microscope (Olympus BX51) using the Komet 6 software. Two technical and four biological replicates were analyzed per condition.

#### Immunofluorescence

Cells were seeded on 8-well Lab-Tek^®^ II Chambered Coverglass plates (Thermo Scientific, 155409) under standard cell culture conditions. Samples were fixed with ice-cold 4% paraformaldehyde in PBS without Ca^2+^ and Mg^2+^. After removing the PFA, the cells were incubated for 20 minutes at room temperature with freshly prepared permeabilization buffer (0.1 % Triton X-100, PBS, pH 7.4). After that, cells were washed in PBS three times for 5 minutes each time.

For cell staining, samples were first rinsed for one hour with blocking buffer (3% fetal bovine serum, PBS, pH 7.4) at room temperature. Then, primary antibodies (OAT, Invitrogen, PA5-66715, 1:500 v/v; Fumarase, Invitrogen #PA5-82899, 1:500 v/v; Casein Kinase 2 beta, Abcam #ab151784, 1:500 v/v; PDHA1 [9H9AF5], Abcam #ab110330, 1:200 v/v, and COX4I1 Abcam #ab33985, 1:500 v/v) were diluted (as indicated above) in blocking buffer and incubated overnight at 4°C. Next, the chambered coverglass plates were rinsed three times with PBS and cells were labeled with fluorescently conjugated secondary antibody (anti-Rabbit Alexa Fluor 488, Invitrogen #A-11008, 1:500 v/v; anti-Mouse Alexa Fluor 647, Abcam # ab150119, 1:500 v/v) in blocking buffer for one hour at room temperature. Afterward, unbound secondary antibodies were removed by washing with PBS three times for 5 minutes each at room temperature. Finally, nuclei were labeled with 1 μg/mL Hoechst 33342 in PBS (Invitrogen #H1399). Imaging was performed in PBS. A Zeiss LSM 710 laser scanning confocal microscope was used, and cells were imaged using x63 oil immersion objective (Plan Apochromat NA 1.40). The number of biological replicates is indicated in the legends of the Figures 5, 6 and 7.

#### Respirometry

One day before the experiment, on four different days, 60,000 cells were plated on XF24 cell plates (Agilent) to measure cell respiration. After irradiation, PBS was replaced by DMEM without sodium bicarbonate, and cells were incubated for 1 hour at 37°C and atmospheric pressure of CO_2_. Oxygen consumption rate (OCR) was measured in a Seahorse XF24 Analyzer (Agilent), before and after subsequent additions of 1 μM oligomycin, 1 μM CCCP and a mix of 1 μM antimycin and 1 μM rotenone. Each compound was added after three cycles of measurements of 3 minutes each. The concentration of CCCP was determined through the previous titration. At the end of the experiments, each well was washed once with PBS and proteins were resuspended in 100 μM ammonium bicarbonate, containing 8 M urea and 1% sodium deoxycholate. After homogenization, protein concentration was determined by using a BCA assay kit. The OCR values were normalized by the amount of total protein in each well. The number of biological replicates is described in the legend of Figure 8.

### QUANTIFICATION AND STATISTICAL ANALYSIS

#### Pre-processing of the proteomics datasets

Raw files of all proteomics experiments performed in this study were processed using MaxQuant (Cox and Mann, 2008). The Andromeda algorithm (Cox et al., 2011) was used for protein identification against the homo sapiens Uniprot database (downloaded August, 2019; 20416 entries). Error mass tolerance for precursors and fragments were set to 4.5 ppm and 20 ppm, respectively. Cysteine carbamidomethylation was selected as a fixed modification and methionine oxidation and *N*-terminal acetylation were selected as variable modifications. Trypsin was set as digestion enzyme, with a maximum of 2 missed cleavages allowed. A maximum FDR of 1% was allowed both for peptides and proteins identification, and for proteins it was calculated using a decoy database created from the reverse ordination of the protein sequences in the Uniprot database. Protein abundances were quantified by the LFQ algorithm (Cox et al., 2014), based on the normalized chromatographic peak integrations calculated by MaxQuant. Other parameters were kept as default.

For the subcellular proteomics assay, each fraction was considered a different sample in the experimental design annotation file required for the MaxQuant analysis. A matrix of relative quantification data (LFQ) (Cox et al., 2014) for proteins in each fraction was obtained and used for subsequent analysis (n = 5 biological replicates). Each protein was normalized by the total sum of the LFQs for a given replicate/cell map, yielding a value between 0 and 1. Proteins that were not quantified in both biological conditions and in at least 45 of the 90 samples were filtered out to remove uninformative fractionation profiles with missing values generated by stochastic fragmentation in the shotgun proteomics approach. Lists of proteins exclusively present in one biological condition are provided in **Supplementary Spreadsheet S8**. We imputed the missing values in the following manner: proteins with valid values in 4 out of 5 biological replicates in a given fraction had their missing values imputed as the average of the 4 valid values, since these values were more likely to be missing at random. The remaning missing values, which were more likely to be values missing not at random due to lack of reproducibility in quantification across replicates, were imputed as the minimum value of the sample. Other methods of imputation were tested, but this was the one that introduced less bias in our analysis (**Figure S1**).

For the total proteome analysis, the data was log-transformed, and matches to the contaminants and reverse database, as well as proteins identified only by modified sites, and missing values (< 30% in all samples) were filtered out. Statistical significance was assessed using a two-tailed Student’s T-test in the Perseus software (Tyanova et al., 2016) with a permutation-based false discovery rate (FDR) of 5% and a S0 parameter of 0.1. The plots were displayed in the R statistical computing environment, using standard libraries and pheatmap. Compartment-specific proteomic analysis of the log_2_ fold changes of irradiated HaCaT cells relative to controls was performed according to (Parca et al., 2018). Briefly, proteins were assigned to compartments according to GO-terms and the fold-changes associated to each compartment were tested for significant difference against the whole proteome (Wilcoxon rank sum test with 0.5% FDR correction). The number of biological replicates can be found in the legend of Figure 8. The asterisks in Figure 8 indicate compartments with corrected p-value lower than 0.05.

#### Dimensionality reduction

Dimensionality reduction was achieved using the t-distributed stochastic neighbor embedding technique (t-SNE) (Maaten and Hinton, 2008). The fractionation data was plotted with different perplexity parameters (perplexity = 30 yielded the best cluster separation). The plots were colored according to categorical subcellular classifications from the Cell Atlas initiative (Rozenblatt-Rosen et al., 2017), Uniprot (The UniProt Consortium, 2017) and Gene Ontology (Ashburner et al., 2000) databases, providing information on the clusterization of different subcellular compartments. T-distributed stochastic neighbor embedding technique was performed using the Rtsne() function from the Rtsne package in the R environment. Results were plotted with the ggplot2 R package (Wickham, 2009) for 2D visualization and with the 3dplot() function from the rgl R package for 3D visualization. All enrichment analysis were performed using g:Profiler (Raudvere et al., 2019), under default configurations.

#### Curation of organellar markers

Organellar markers were selected based on the curation of proteins classified as pertaining exclusively to one of the following subcellular compartments: cytosol, nucleus, mitochondria and secretory organelles, according to Uniprot and GO classifications. We also incorporated in the marker set a few organellar markers from a previous subcellular proteomics study (Geladaki et al., 2019). Finally, markers also had to be present in all replicates of the experiment. Organellar markers from four different compartments were assigned with different colors to visualize clusterization in the t-SNE plots. The secretory compartment comprises proteins initially assigned to peroxisomes, endoplasmic reticulum, plasma membrane, Golgi apparatus and lysosomes. These organelles were grouped under the term “secretory” because they share similar fractionation profiles that were not well distinguished by the machine learning algorithms in preliminary analysis we performed.

#### Neural networks predictions

A supervised machine learning approach was used for the subcellular localization prediction (Breckels et al., 2018; Gatto et al., 2014b). We used a model of an averaged neural networks algorithm (Gatto et al., 2014b) implemented in the R package pRoloc to produce the paper’s results, but a support vector machine was also tested and yielded similar results. The organellar markers were used to train the model for subcellular localization prediction. Organellar markers were divided into a training and validation set (80/20% proportion for each set) with a 5-fold cross-validation through 100 iterations of the algorithm. We used a grid search to achieve hyperparameter tuning. The accuracy of the classifier was estimated through the F1 score (Breckels et al., 2018), and the best hyperparameters were chosen according to the accuracy of the classifier. The best network size decay for each replicate is described in detail in **Figure S3M**. To filter low prediction scores, we adopted the method performed by Beltran et al., 2016. Briefly, we generated a null distribution of prediction scores through randomization of the empirical scores over 1000 iterations. Then, we estimated an empirical cumulative function from the null distribution using the R function ecdf(). Finally, we set thresholds per organelle and per sample at a cumulative probability of 0.95.

To define a translocation, we performed the same neural network analysis on the whole set of controls and on the whole set of irradiated samples, in order to obtain a consensus classification for each biological condition. The low scores were filtered according to the same principle described above. F1 scores obtained during optimization of the hyperparameters are shown in **Figure S3**.

To compare subcellular localizations classified by the learning algorithm to GO, we used the biomaRt (Durinck et al., 2005) and GO.db R packages.

#### TRANSPIRE analysis

The TRANSPIRE pipeline, developed by Kennedy et al., 2020, was used for the translocation prediction. Curated organellar markers were utilized to generate synthetic translocations, which are then used to train the learning algorithm in distinguishing translocation classes and consequently translocating from non-translocating proteins. In brief, each organellar marker in the control samples is concatenated with every other organellar marker of the treated samples, producing synthetic translocations and non-translocations (when markers of the same compartment are concatenated). For example, a synthetic translocation that simulates the migration of a protein from the nucleus to the cytosol would have a fractionation profile that is characterized by the combination of a nuclear marker profile in all control samples (45 fractions) with the cytosolic marker profile in the treated samples (also 45 fractions, yielding a total of 90 “fractions” per synthetic translocation).

Synthetic translocations were used to train a Stochastic Variational Gaussian Process Classifier (SVGPC) implemented in TRANSPIRE through the GPFlow package (built upon the TensorFlow platform in Python). This model is composed of a kernel function, a likelihood function, n latent variables (which account for the number of translocation classes), a training set, and a subset of the training set used as inducing points. The model implemented in TRANSPIRE uses softmax as a likelihood function to improve score calibration.

Hyperparameter tuning involved choosing the kernel type (squared exponential, rational quadratic, exponential Matern32 and Matern52, as implemented by TRANSPIRE through GPFlow) and the number of inducing points (ranging from 1 to 500). The synthetic translocation data were divided into training, validation, and test sets in a 50/20/20% proportion, respectively, during training. The training data was further split into five balanced folds during hyperparameter tuning, allowing for a 5-fold cross-validation. A class imbalance was prevented by allowing the most frequent translocation classes to have, at most, three times more proteins than the least frequent. The best hyperparameters selected through the grid search were the squared exponential kernel and 30 inducing points (optimization plots are shown in the Supporting Information). The results were evaluated by maximizing the evidence lower bound (ELBO) using the Adam optimizer. Afterward, the resulting model was used to predict translocations in the actual dataset and performance was evaluated based on the held-out test partition of the synthetic translocation data.

The output of the TRANSPIRE pipeline entails the classification of a translocation class (e.g., “Nucleus to Cytosol”) for each protein plus a classifier score. The classifier score ranges from 0 to 1 for each translocation class, and the sum of the scores for all classes for each protein should be equal to 1. Class prediction is based on the highest classifier score for a given translocation class. This score is referred to as “predicted scores” in the spreadsheets in the Supporting Information. Additionally, the TRANSPIRE pipeline provides a translocation score, defined as the sum of the predicted scores for all true translocation classes. This score accounts for situations in which high classifier scores are split among at least two translocation classes.

TRANSPIRE also allows for the computation of false-positive rates (FPR), based on the model’s performance, setting thresholds for the translocation scores to minimize the likelihood of false positives. Herein, we adopted a 0.1% FPR to generate a more stringent list of translocation targets. The entire pipeline was performed as described by Kennedy et al., 2020, in Python version 3.7.6.

#### Spreadsheets Titles

**Spreadsheet S1 – Total label-free quantification. Related to Figures 1 and 2.**

Non-filtered and non-imputed label-free quantification of proteins across organelle fractions from the differential centrifugation.

**Spreadsheet S2 – Filtered and imputed label-free quantification. Related to Figures 1 and 2.**

Filtered and imputed label-free quantification of proteins across organelle fractions

**Spreadsheet S3 – Organellar markers. Related to Figures 2 and 3.**

List of refined organellar markers used to predict subcellular localization and translocations

**Spreadsheet S4 – Neural networks classifications. Related to Figure 2.**

Neural networks predictions of subcellular localization along with the respective prediction scores

**Spreadsheet S5 – Translocation predictions. Related to Figure 3.**

Translocation events predicted by the TRANSPIRE algorithm in HaCaT cells upon UVA exposure

**Spreadsheet S6 – Comparison of neural networks and TRANSPIRE results. Related to Figure 4.**

Comparison of the classifications obtained from the TRANSPIRE and neural network algorithm (tab 1) and enrichment analysis of protein set (tab 2)

**Spreadsheet S7 – Label-free quantification of whole-cell lysates. Related to Figure 8.**

Label-free quantification data and statistical analysis of whole-cell lysates obtained from irradiated and control HaCaT cells 1h30 after exposure to UVA

**Spreadsheet S8 – Proteins exclusively present in one condition. Related to Figure 8.**

Proteins exclusively present in one biological condition in the spatial proteomics dataset

## References

Andersen, J.S., Wilkinson, C.J., Mayor, T., Mortensen, P., Nigg, E.A., Mann, M., 2003. Proteomic characterization of the human centrosome by protein correlation profiling. Nature 426, 570–574. https://doi.org/10.1038/nature02166

Ashburner, M., Ball, C.A., Blake, J.A., Botstein, D., Butler, H., Cherry, J.M., Davis, A.P., Dolinski, K., Dwight, S.S., Eppig, J.T., Harris, M.A., Hill, D.P., Issel-Tarver, L., Kasarskis, A., Lewis, S., Matese, J.C., Richardson, J.E., Ringwald, M., Rubin, G.M., Sherlock, G., 2000. Gene Ontology: tool for the unification of biology. Nature Genetics 25, 25–29. https://doi.org/10.1038/75556

Berneburg, M., Plettenberg, H., Medve-König, K., Pfahlberg, A., Gers-Barlag, H., Gefeller, O., Krutmann, J., 2004. Induction of the Photoaging-Associated Mitochondrial Common Deletion In Vivo in Normal Human Skin. J Invest Dermatol 122, 1277–1283. https://doi.org/10.1111/j.0022-202X.2004.22502.x

Boukamp, P., Bleuel, K., Popp, S., Vormwald-Dogan, V., Fusenig, N.E., 1997. Functional evidence for tumor-suppressor activity on chromosome 15 in human skin carcinoma cells and thrombospondin-1 as the potential suppressor. Journal of Cellular Physiology 173, 256–260. https://doi.org/10.1002/(SICI)1097-4652(199711)173:2<256::AID-JCP31>3.0.CO;2-B

Breckels, L.M., Mulvey, C.M., Lilley, K.S., Gatto, L., 2018. A Bioconductor workflow for processing and analysing spatial proteomics data. F1000Res 5, 2926. https://doi.org/10.12688/f1000research.10411.2

Breker, M., Schuldiner, M., 2014. The emergence of proteome-wide technologies: systematic analysis of proteins comes of age. Nature Reviews Molecular Cell Biology 15, 453–464. https://doi.org/10.1038/nrm3821

Chatzidoukaki, O., Goulielmaki, E., Schumacher, B., Garinis, G.A., 2020. DNA Damage Response and Metabolic Reprogramming in Health and Disease. Trends in Genetics 36, 777–791. https://doi.org/10.1016/j.tig.2020.06.018

Chen, H., Weng, Q.Y., Fisher, D.E., 2014. UV Signaling Pathways within the Skin. J Invest Dermatol 134, 2080–2085. https://doi.org/10.1038/jid.2014.161

Cochet, C., Chambaz, E.M., 1983. Oligomeric structure and catalytic activity of G type casein kinase. Isolation of the two subunits and renaturation experiments. J Biol Chem 258, 1403–1406.

Coin, F., Auriol, J., Tapias, A., Clivio, P., Vermeulen, W., Egly, J.-M., 2004. Phosphorylation of XPB helicase regulates TFIIH nucleotide excision repair activity. The EMBO Journal 23, 4835–4846. https://doi.org/10.1038/sj.emboj.7600480

Cox, J., Hein, M.Y., Luber, C.A., Paron, I., Nagaraj, N., Mann, M., 2014. Accurate Proteome-wide Label-free Quantification by Delayed Normalization and Maximal Peptide Ratio Extraction, Termed MaxLFQ *. Molecular & Cellular Proteomics 13, 2513–2526. https://doi.org/10.1074/mcp.M113.031591

Cox, J., Mann, M., 2008. MaxQuant enables high peptide identification rates, individualized p.p.b.-range mass accuracies and proteome-wide protein quantification. Nature Biotechnology 26, 1367–1372. https://doi.org/10.1038/nbt.1511

Cox, J., Neuhauser, N., Michalski, A., Scheltema, R.A., Olsen, J.V., Mann, M., 2011. Andromeda: A Peptide Search Engine Integrated into the MaxQuant Environment. J. Proteome Res. 10, 1794–1805. https://doi.org/10.1021/pr101065j

Dabke, K., Kreimer, S., Jones, M.R., Parker, S.J., 2021. A Simple Optimization Workflow to Enable Precise and Accurate Imputation of Missing Values in Proteomic Data Sets. Journal of Proteome Research. https://doi.org/10.1021/acs.jproteome.1c00070

de Gruijl, F.R., van Kranen, H.J., Mullenders, L.H.F., 2001. UV-induced DNA damage, repair, mutations and oncogenic pathways in skin cancer. Journal of Photochemistry and Photobiology B: Biology, Consequences of exposure to sunlight:elements to assess protection 63, 19–27. https://doi.org/10.1016/S1011-1344(01)00199-3

Delinasios, G.J., Karbaschi, M., Cooke, M.S., Young, A.R., 2018. Vitamin E inhibits the UVAI induction of “light” and “dark” cyclobutane pyrimidine dimers, and oxidatively generated DNA damage, in keratinocytes. Sci Rep 8, 1–12. https://doi.org/10.1038/s41598-017-18924-4

Djavaheri-Mergny, M., Marsac, C., Mazière, C., Santus, R., Michel, L., Dubertret, L., Mazière, J.C., 2001. UV-A irradiation induces a decrease in the mitochondrial respiratory activity of human NCTC 2544 keratinocytes. Free Radical Research 34, 583–594. https://doi.org/10.1080/10715760100300481

Durinck, S., Moreau, Y., Kasprzyk, A., Davis, S., De Moor, B., Brazma, A., Huber, W., 2005. BioMart and Bioconductor: a powerful link between biological databases and microarray data analysis. Bioinformatics 21, 3439–3440. https://doi.org/10.1093/bioinformatics/bti525

Edifizi, D., Nolte, H., Babu, V., Castells-Roca, L., Mueller, M.M., Brodesser, S., Krüger, M., Schumacher, B., 2017. Multilayered Reprogramming in Response to Persistent DNA Damage in C. elegans. Cell Reports 20, 2026–2043. https://doi.org/10.1016/j.celrep.2017.08.028

El Ghissassi, F., Baan, R., Straif, K., Grosse, Y., Secretan, B., Bouvard, V., Benbrahim-Tallaa, L., Guha, N., Freeman, C., Galichet, L., Cogliano, V., WHO International Agency for Research on Cancer Monograph Working Group, 2009. A review of human carcinogens--part D: radiation. Lancet Oncol. 10, 751–752. https://doi.org/10.1016/s1470-2045(09)70213-x

Elia, A.E.H., Boardman, A.P., Wang, D.C., Huttlin, E.L., Everley, R.A., Dephoure, N., Zhou, C., Koren, I., Gygi, S.P., Elledge, S.J., 2015. Quantitative Proteomic Atlas of Ubiquitination and Acetylation in the DNA Damage Response. Mol. Cell 59, 867–881. https://doi.org/10.1016/j.molcel.2015.05.006

Filhol, O., Baudier, J., Delphin, C., Loue-Mackenbach, P., Chambaz, E.M., Cochet, C., 1992. Casein kinase II and the tumor suppressor protein P53 associate in a molecular complex that is negatively regulated upon P53 phosphorylation. J. Biol. Chem. 267, 20577–20583.

Foster, L.J., Hoog, C.L. de, Zhang, Yanling, Zhang, Yong, Xie, X., Mootha, V.K., Mann, M., 2006. A Mammalian Organelle Map by Protein Correlation Profiling. Cell 125, 187–199. https://doi.org/10.1016/j.cell.2006.03.022

Fusenig, N.E., Boukamp, P., 1998. Multiple stages and genetic alterations in immortalization, malignant transformation, and tumor progression of human skin keratinocytes. Molecular Carcinogenesis 23, 144–158. https://doi.org/10.1002/(SICI)1098-2744(199811)23:3<144::AID-MC3>3.0.CO;2-U

Gatto, L., Breckels, L.M., Burger, T., Nightingale, D.J.H., Groen, A.J., Campbell, C., Nikolovski, N., Mulvey, C.M., Christoforou, A., Ferro, M., Lilley, K.S., 2014a. A Foundation for Reliable Spatial Proteomics Data Analysis. Mol Cell Proteomics 13, 1937–1952. https://doi.org/10.1074/mcp.M113.036350

Gatto, L., Breckels, L.M., Wieczorek, S., Burger, T., Lilley, K.S., 2014b. Mass-spectrometry-based spatial proteomics data analysis using pRoloc and pRolocdata. Bioinformatics 30, 1322–1324. https://doi.org/10.1093/bioinformatics/btu013

Geladaki, A., Kočevar Britovšek, N., Breckels, L.M., Smith, T.S., Vennard, O.L., Mulvey, C.M., Crook, O.M., Gatto, L., Lilley, K.S., 2019. Combining LOPIT with differential ultracentrifugation for high-resolution spatial proteomics. Nature Communications 10, 331. https://doi.org/10.1038/s41467-018-08191-w

Gray, G.K., McFarland, B.C., Rowse, A.L., Gibson, S.A., Benveniste, E.N., 2014. Therapeutic CK2 inhibition attenuates diverse prosurvival signaling cascades and decreases cell viability in human breast cancer cells. Oncotarget 5, 6484–6496. https://doi.org/10.18632/oncotarget.2248

Grecu, D., Assairi, L., 2014. CK2 phosphorylation of human centrins 1 and 2 regulates their binding to the DNA repair protein XPC, the centrosomal protein Sfi1 and the phototransduction protein transducin β. FEBS Open Bio 4, 407–419. https://doi.org/10.1016/j.fob.2014.04.002

Halliday, G.M., Rana, S., 2008. Waveband and Dose Dependency of Sunlight-induced Immunomodulation and Cellular Changes†. Photochemistry and Photobiology 84, 35–46. https://doi.org/10.1111/j.1751-1097.2007.00212.x

He, Y.-Y., Huang, J.-L., Sik, R.H., Chignell, C.F., Liu, J., Waalkes, M.P., 2004. Expression Profiling of Human Keratinocyte Response to Ultraviolet A: Implications in Apoptosis. Journal of Investigative Dermatology 122, 533–543. https://doi.org/10.1046/j.0022-202X.2003.22123.x

Ikehata, H., 2018. Mechanistic considerations on the wavelength-dependent variations of UVR genotoxicity and mutagenesis in skin: The discrimination of UVA-signature from UV-signature mutation. Photochem Photobiol Sci 17, 1861–1871. https://doi.org/10.1039/c7pp00360a

Itzhak, D.N., Davies, C., Tyanova, S., Mishra, A., Williamson, J., Antrobus, R., Cox, J., Weekes, M.P., Borner, G.H.H., 2017. A Mass Spectrometry-Based Approach for Mapping Protein Subcellular Localization Reveals the Spatial Proteome of Mouse Primary Neurons. Cell Reports 20, 2706–2718. https://doi.org/10.1016/j.celrep.2017.08.063

Jean Beltran, P.M., Mathias, R.A., Cristea, I.M., 2016. A Portrait of the Human Organelle Proteome In Space and Time during Cytomegalovirus Infection. cels 3, 361–373.e6. https://doi.org/10.1016/j.cels.2016.08.012

Jugé, R., Breugnot, J., Da Silva, C., Bordes, S., Closs, B., Aouacheria, A., 2016. Quantification and Characterization of UVB-Induced Mitochondrial Fragmentation in Normal Primary Human Keratinocytes. Sci Rep 6, 35065. https://doi.org/10.1038/srep35065

Kennedy, M.A., Hofstadter, W.A., Cristea, I.M., 2020. TRANSPIRE: A Computational Pipeline to Elucidate Intracellular Protein Movements from Spatial Proteomics Data Sets. J. Am. Soc. Mass Spectrom. 31, 1422–1439. https://doi.org/10.1021/jasms.0c00033

Klein, A.M., Brash, D.E., Jones, P.H., Simons, B.D., 2010. Stochastic fate of p53-mutant epidermal progenitor cells is tilted toward proliferation by UV B during preneoplasia. PNAS 107, 270–275. https://doi.org/10.1073/pnas.0909738107

Kowaltowski, A.J., Menezes-Filho, S.L., Assali, E.A., Gonçalves, I.G., Cabral-Costa, J.V., Abreu, P., Miller, N., Nolasco, P., Laurindo, F.R.M., Bruni-Cardoso, A., Shirihai, O.S., 2019. Mitochondrial morphology regulates organellar Ca2+ uptake and changes cellular Ca2+ homeostasis. The FASEB Journal 33, 13176–13188. https://doi.org/10.1096/fj.201901136R

Krahmer, N., Najafi, B., Schueder, F., Quagliarini, F., Steger, M., Seitz, S., Kasper, R., Salinas, F., Cox, J., Uhlenhaut, N.H., Walther, T.C., Jungmann, R., Zeigerer, A., Borner, G.H.H., Mann, M., 2018. Organellar Proteomics and Phospho-Proteomics Reveal Subcellular Reorganization in Diet-Induced Hepatic Steatosis. Developmental Cell 47, 205–221.e7. https://doi.org/10.1016/j.devcel.2018.09.017

Larance, M., Lamond, A.I., 2015. Multidimensional proteomics for cell biology. Nature Reviews Molecular Cell Biology 16, 269–280. https://doi.org/10.1038/nrm3970

Lehman, T.A., Modali, R., Boukamp, P., Stanek, J., Bennett, W.P., Welsh, J.A., Metcalf, R.A., Stampfer, M.R., Fusenig, N., Rogan, E.M., Harris, C.C., 1993. p53 Mutations in human immortalized epithelial cell lines. Carcinogenesis 14, 833–839. https://doi.org/10.1093/carcin/14.5.833

Liu, M., Dongre, A., 2021. Proper imputation of missing values in proteomics datasets for differential expression analysis. Briefings in Bioinformatics 22, bbaa112. https://doi.org/10.1093/bib/bbaa112

Lundberg, E., Borner, G.H.H., 2019. Spatial proteomics: a powerful discovery tool for cell biology. Nat. Rev. Mol. Cell Biol. 20, 285–302. https://doi.org/10.1038/s41580-018-0094-y

Maaten, L. van der Hinton, G., 2008. Visualizing Data using t-SNE. Journal of Machine Learning Research 9, 2579–2605.

Maresca, V., Flori, E., Briganti, S., Camera, E., Cario-André, M., Taïeb, A., Picardo, M., 2006. UVA-Induced Modification of Catalase Charge Properties in the Epidermis Is Correlated with the Skin Phototype. Journal of Investigative Dermatology 126, 182–190. https://doi.org/10.1038/sj.jid.5700021

Mattiazzi Usaj, M., Styles, E.B., Verster, A.J., Friesen, H., Boone, C., Andrews, B.J., 2016. High-Content Screening for Quantitative Cell Biology. Trends Cell Biol. 26, 598–611. https://doi.org/10.1016/j.tcb.2016.03.008

Montenarh, M., 2016. Protein kinase CK2 in DNA damage and repair. Translational Cancer Research 5. https://doi.org/10.21037/6469

Moreno, N.C., de Souza, T.A., Garcia, C.C.M., Ruiz, N.Q., Corradi, C., Castro, L.P., Munford, V., Ienne, S., Alexandrov, L.B., Menck, C.F.M., 2020. Whole-exome sequencing reveals the impact of UVA light mutagenesis in xeroderma pigmentosum variant human cells. Nucleic Acids Research 48, 1941–1953. https://doi.org/10.1093/nar/gkz1182

Mueller, M.M., Peter, W., Mappes, M., Huelsen, A., Steinbauer, H., Boukamp, P., Vaccariello, M., Garlick, J., Fusenig, N.E., 2001. Tumor Progression of Skin Carcinoma Cells in Vivo Promoted by Clonal Selection, Mutagenesis, and Autocrine Growth Regulation by Granulocyte Colony-Stimulating Factor and Granulocyte-Macrophage Colony-Stimulating Factor. Am J Pathol 159, 1567–1579.

Mulvey, C.M., Breckels, L.M., Geladaki, A., Britovšek, N.K., Nightingale, D.J.H., Christoforou, A., Elzek, M., Deery, M.J., Gatto, L., Lilley, K.S., 2017. Using hyperLOPIT to perform high-resolution mapping of the spatial proteome. Nat Protoc 12, 1110–1135. https://doi.org/10.1038/nprot.2017.026

Palla, G., Fischer, D.S., Regev, A., Theis, F.J., 2022. Spatial components of molecular tissue biology. Nat Biotechnol 1–11. https://doi.org/10.1038/s41587-021-01182-1

Parca, L., Beck, M., Bork, P., Ori, A., 2018. Quantifying compartment-associated variations of protein abundance in proteomics data. Molecular Systems Biology 14. https://doi.org/10.15252/msb.20178131

Pouli, D., Balu, M., Alonzo, C.A., Liu, Z., Quinn, K.P., Rius-Diaz, F., Harris, R.M., Kelly, K.M., Tromberg, B.J., Georgakoudi, I., 2016. Imaging mitochondrial dynamics in human skin reveals depth-dependent hypoxia and malignant potential for diagnosis. Science Translational Medicine 8, 367ra169–367ra169. https://doi.org/10.1126/scitranslmed.aag2202

Premi, S., Wallisch, S., Mano, C.M., Weiner, A.B., Bacchiocchi, A., Wakamatsu, K., Bechara, E.J.H., Halaban, R., Douki, T., Brash, D.E., 2015. Chemiexcitation of melanin derivatives induces DNA photoproducts long after UV exposure. Science 347, 842–847. https://doi.org/10.1126/science.1256022

Rappsilber, J., Mann, M., Ishihama, Y., 2007. Protocol for micro-purification, enrichment, pre-fractionation and storage of peptides for proteomics using StageTips. Nat Protoc 2, 1896–1906. https://doi.org/10.1038/nprot.2007.261

Raudvere, U., Kolberg, L., Kuzmin, I., Arak, T., Adler, P., Peterson, H., Vilo, J., 2019. g:Profiler: a web server for functional enrichment analysis and conversions of gene lists (2019 update). Nucleic Acids Research 47, W191–W198. https://doi.org/10.1093/nar/gkz369

Ren, Q., Kari, C., Quadros, M.R.D., Burd, R., McCue, P., Dicker, A.P., Rodeck, U., 2006. Malignant Transformation of Immortalized HaCaT Keratinocytes through Deregulated Nuclear Factor κB Signaling. Cancer Res 66, 5209–5215. https://doi.org/10.1158/0008-5472.CAN-05-4158

Rozenblatt-Rosen, O., Stubbington, M.J.T., Regev, A., Teichmann, S.A., 2017. The Human Cell Atlas: from vision to reality. Nature 550, 451–453. https://doi.org/10.1038/550451a

Sabouny, R., Shutt, T.E., 2020. Reciprocal Regulation of Mitochondrial Fission and Fusion. Trends in Biochemical Sciences 45, 564–577. https://doi.org/10.1016/j.tibs.2020.03.009

Schuch, A.P., Moreno, N.C., Schuch, N.J., Menck, C.F.M., Garcia, C.C.M., 2017. Sunlight damage to cellular DNA: Focus on oxidatively generated lesions. Free Radic. Biol. Med. 107, 110–124. https://doi.org/10.1016/j.freeradbiomed.2017.01.029

Simpson, C.L., Tokito, M.K., Uppala, R., Sarkar, M.K., Gudjonsson, J.E., Holzbaur, E.L.F., 2021. NIX initiates mitochondrial fragmentation via DRP1 to drive epidermal differentiation. Cell Reports 34, 108689. https://doi.org/10.1016/j.celrep.2021.108689

Sprenger, H.-G., Langer, T., 2019. The Good and the Bad of Mitochondrial Breakups. Trends in Cell Biology 29, 888–900. https://doi.org/10.1016/j.tcb.2019.08.003

Teitz, T., Eli, D., Penner, M., Bakhanashvili, M., Naiman, T., Timme, T.L., Wood, C.M., Moses, R.E., Canaani, D., 1990. Expression of the cDNA for the beta subunit of human casein kinase II confers partial UV resistance on xeroderma pigmentosum cells. Mutation Research/DNA Repair 236, 85–97. https://doi.org/10.1016/0921-8777(90)90036-5

Thul, P.J., Åkesson, L., Wiking, M., Mahdessian, D., Geladaki, A., Ait Blal, H., Alm, T., Asplund, A., Björk, L., Breckels, L.M., Bäckström, A., Danielsson, F., Fagerberg, L., Fall, J., Gatto, L., Gnann, C., Hober, S., Hjelmare, M., Johansson, F., Lee, S., Lindskog, C., Mulder, J., Mulvey, C.M., Nilsson, P., Oksvold, P., Rockberg, J., Schutten, R., Schwenk, J.M., Sivertsson, Å., Sjöstedt, E., Skogs, M., Stadler, C., Sullivan, D.P., Tegel, H., Winsnes, C., Zhang, C., Zwahlen, M., Mardinoglu, A., Pontén, F., von Feilitzen, K., Lilley, K.S., Uhlén, M., Lundberg, E., 2017. A subcellular map of the human proteome. Science 356. https://doi.org/10.1126/science.aal3321

Tondera, D., Grandemange, S., Jourdain, A., Karbowski, M., Mattenberger, Y., Herzig, S., Da Cruz, S., Clerc, P., Raschke, I., Merkwirth, C., Ehses, S., Krause, F., Chan, D.C., Alexander, C., Bauer, C., Youle, R., Langer, T., Martinou, J.-C., 2009. SLP-2 is required for stress-induced mitochondrial hyperfusion. The EMBO Journal 28, 1589–1600. https://doi.org/10.1038/emboj.2009.89

Tripodi, F., Zinzalla, V., Vanoni, M., Alberghina, L., Coccetti, P., 2007. In CK2 inactivated cells the cyclin dependent kinase inhibitor Sic1 is involved in cell-cycle arrest before the onset of S phase. Biochemical and Biophysical Research Communications 359, 921–927. https://doi.org/10.1016/j.bbrc.2007.05.195

Twig, G., Shirihai, O.S., 2011. The Interplay Between Mitochondrial Dynamics and Mitophagy. Antioxid Redox Signal 14, 1939–1951. https://doi.org/10.1089/ars.2010.3779

Tyanova, S., Temu, T., Sinitcyn, P., Carlson, A., Hein, M.Y., Geiger, T., Mann, M., Cox, J., 2016. The Perseus computational platform for comprehensive analysis of (prote)omics data. Nature Methods 13, 731–740. https://doi.org/10.1038/nmeth.3901

The UniProt Consortium, 2017. UniProt: the universal protein knowledgebase. Nucleic Acids Research 45, D158–D169. https://doi.org/10.1093/nar/gkw1099

Wang, Ping, Wang, Peiguo, Liu, B., Zhao, J., Pang, Q., Agrawal, S.G., Jia, L., Liu, F.-T., 2015. Dynamin-related protein Drp1 is required for Bax translocation to mitochondria in response to irradiation-induced apoptosis. Oncotarget 6, 22598–22612. https://doi.org/10.18632/oncotarget.4200

Wickham, H., 2009. Polishing your plots for publication, in: Wickham, H. (Ed.), Ggplot2: Elegant Graphics for Data Analysis, Use R. Springer, New York, NY, pp. 139–155. https://doi.org/10.1007/978-0-387-98141-3_8

Wondrak, G.T., Jacobson, M.K., Jacobson, E.L., 2006. Endogenous UVA-photosensitizers: mediators of skin photodamage and novel targets for skin photoprotection. Photochem. Photobiol. Sci. 5, 215–237. https://doi.org/10.1039/B504573H

Yefi, R., Ponce, D.P., Niechi, I., Silva, E., Cabello, P., Rodriguez, D.A., Marcelain, K., Armisen, R., Quest, A.F.G., Tapia, J.C., 2011. Protein kinase CK2 promotes cancer cell viability via up-regulation of cyclooxygenase-2 expression and enhanced prostaglandin E2 production. Journal of Cellular Biochemistry 112, 3167–3175. https://doi.org/10.1002/jcb.23247

Zhang, Z., Liu, L., Wu, S., Xing, D., 2016. Drp1, Mff, Fis1, and MiD51 are coordinated to mediate mitochondrial fission during UV irradiation-induced apoptosis. The FASEB Journal 30, 466–476. https://doi.org/10.1096/fj.15-274258

Zhou, C., Elia, A.E.H., Naylor, M.L., Dephoure, N., Ballif, B.A., Goel, G., Xu, Q., Ng, A., Chou, D.M., Xavier, R.J., Gygi, S.P., Elledge, S.J., 2016. Profiling DNA damage-induced phosphorylation in budding yeast reveals diverse signaling networks. Proc Natl Acad Sci USA 113, E3667–E3675. https://doi.org/10.1073/pnas.1602827113

