## Supplemental Figures for "Spatial proteomics reveals subcellular reorganization in human keratinocytes exposed to UVA light"

### A. Imputation = 0

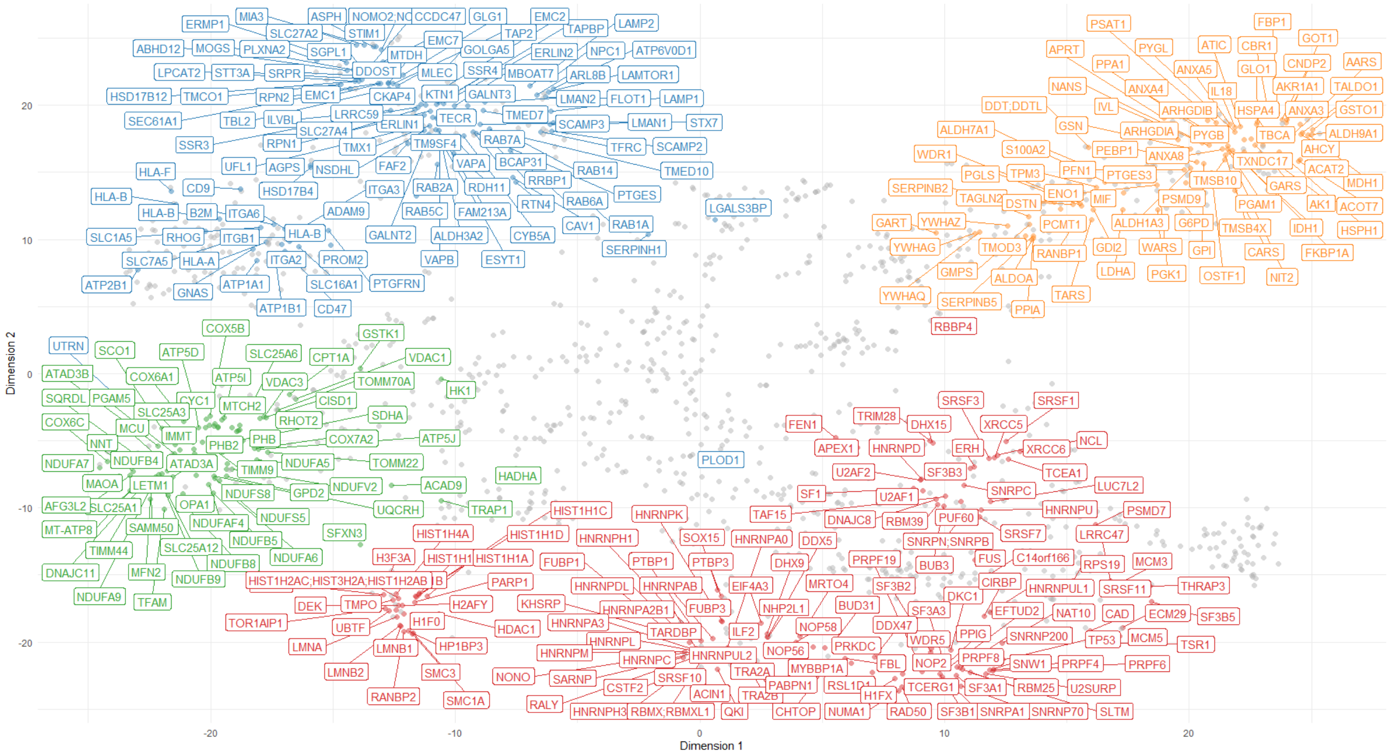


### B. Imputation = bpca (Bayesian PCA Missing Value Estimation)

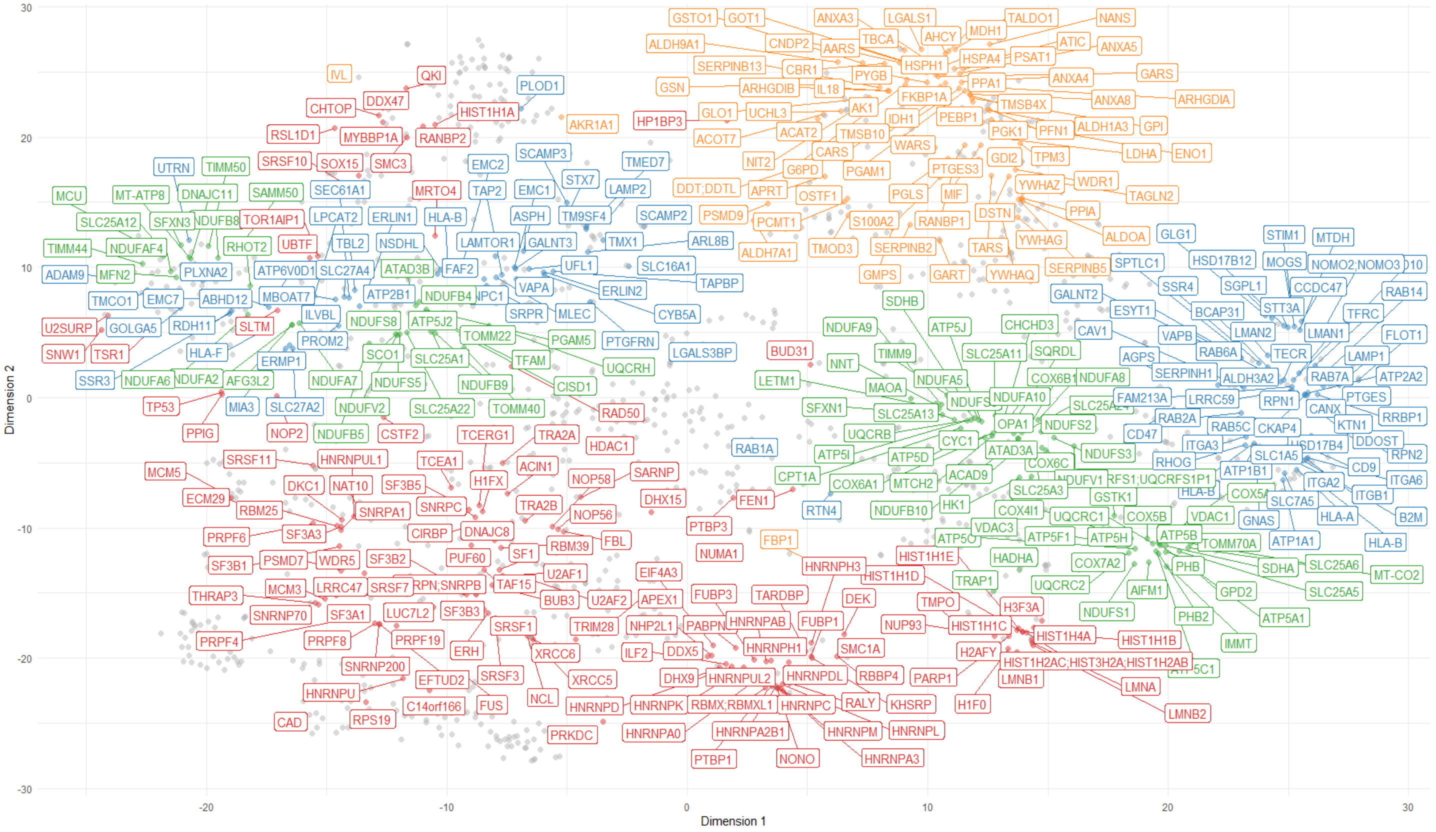

### C. Imputation = QRILC

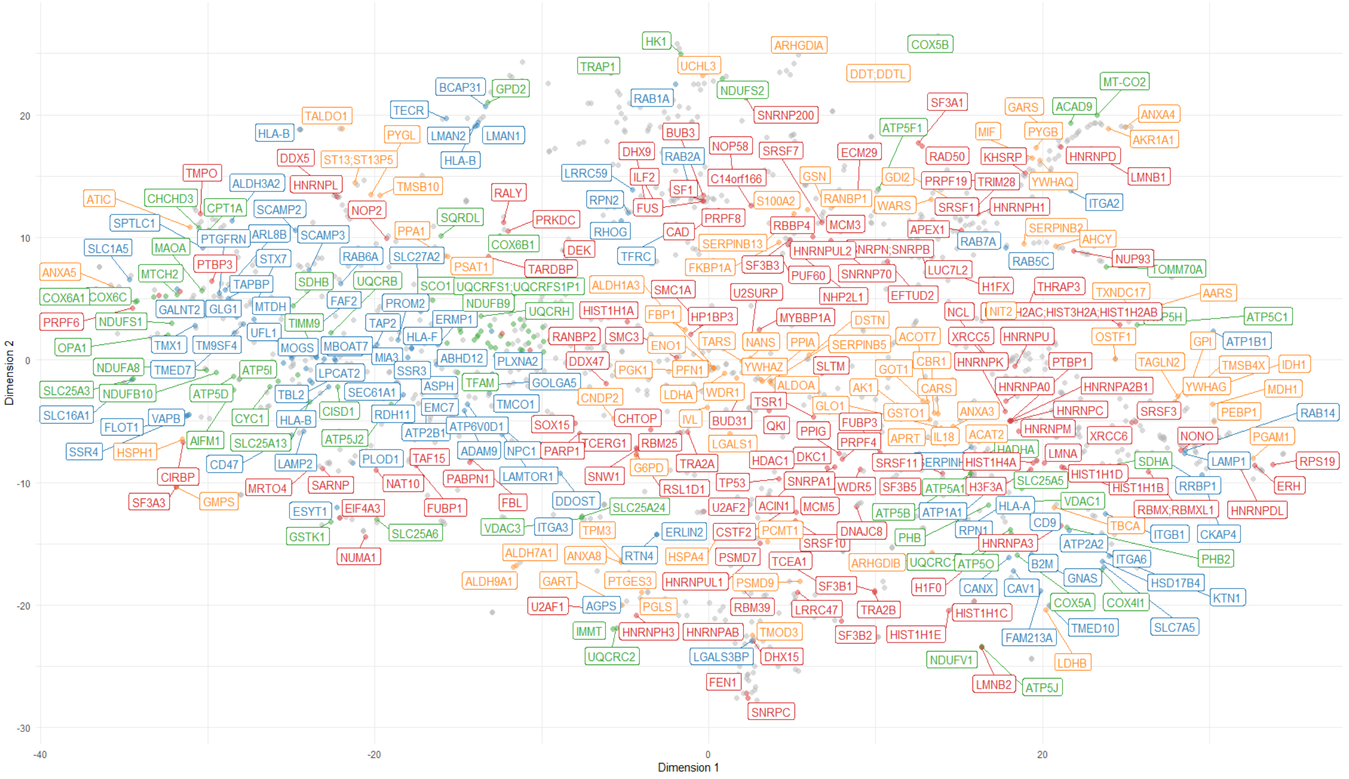


### D. Imputation = For proteins quantified in 4 out of 5 replicates of a given fraction, the missing value was imputed as the average of the valid values. The remaining missing values were imputed as the minimum value of the sample.

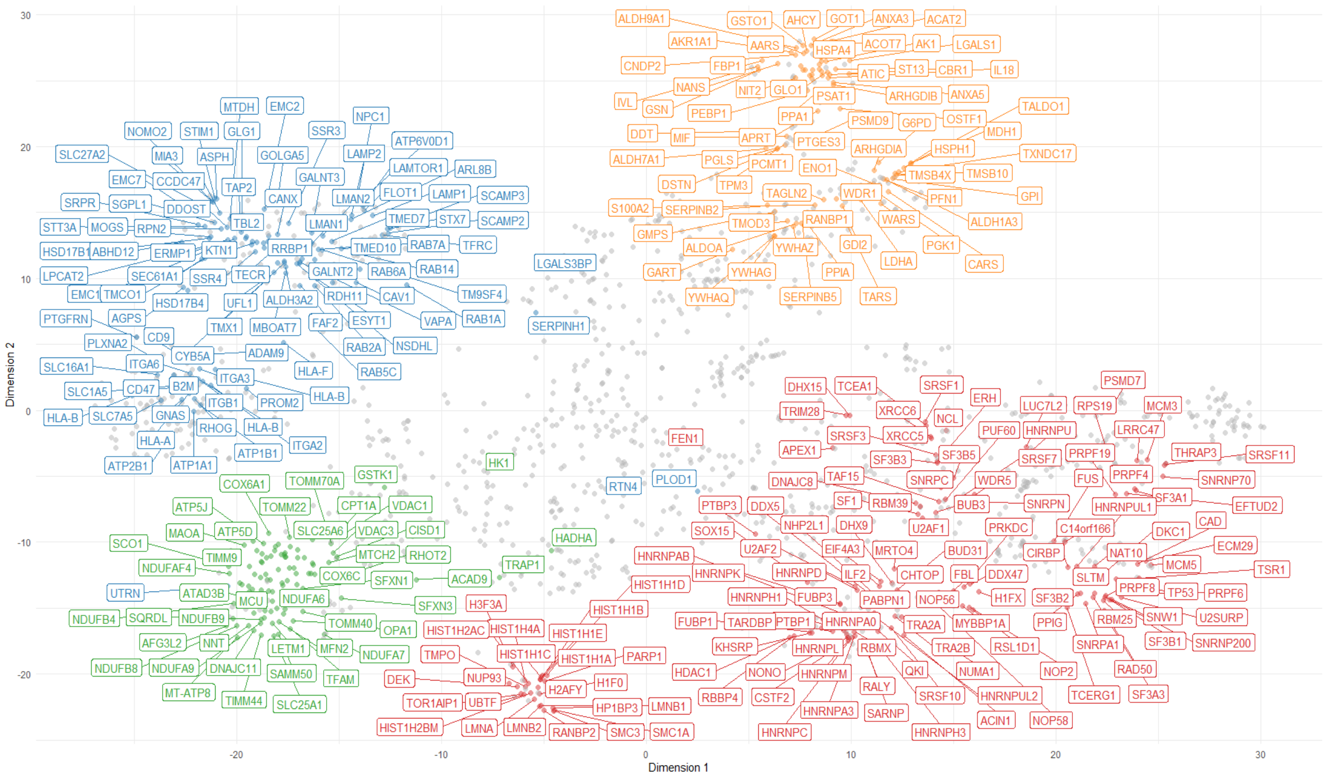

### E. Controls

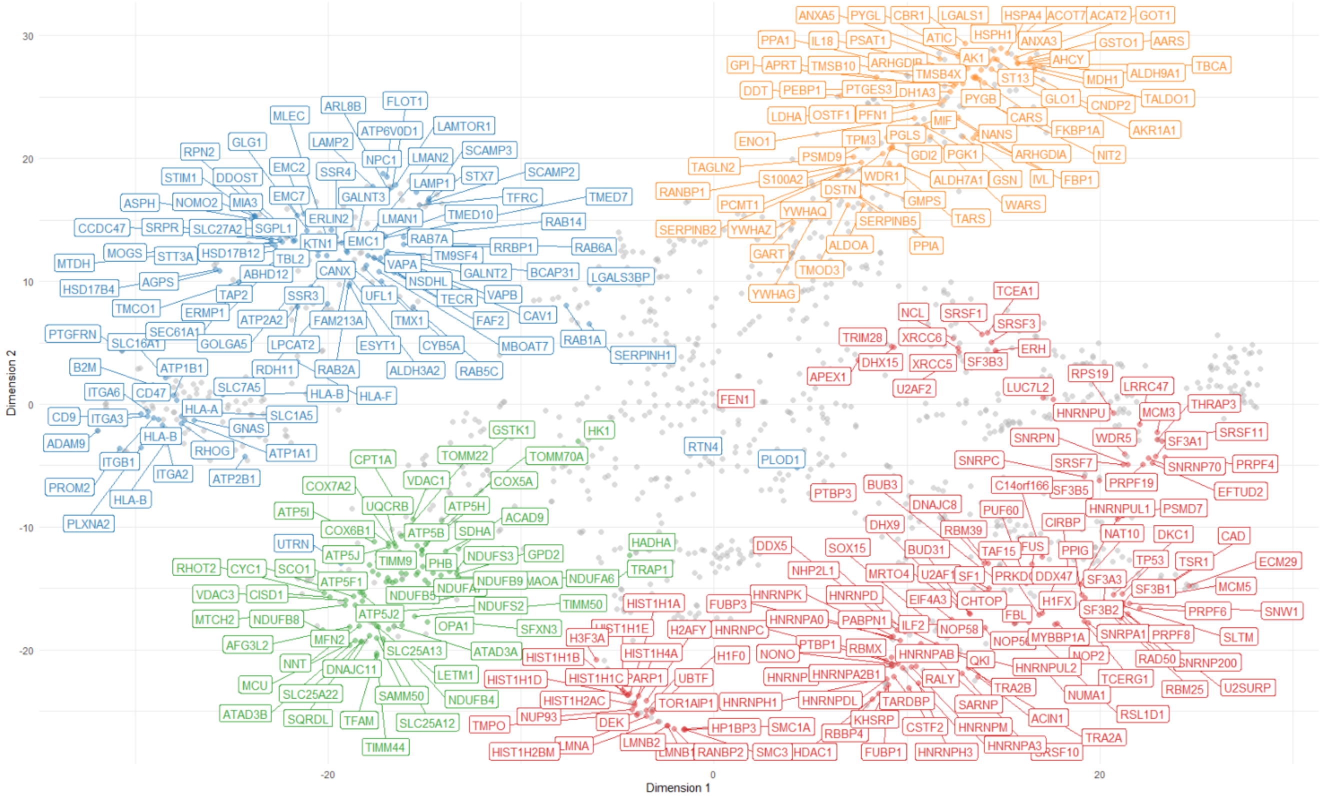


### F. UVA

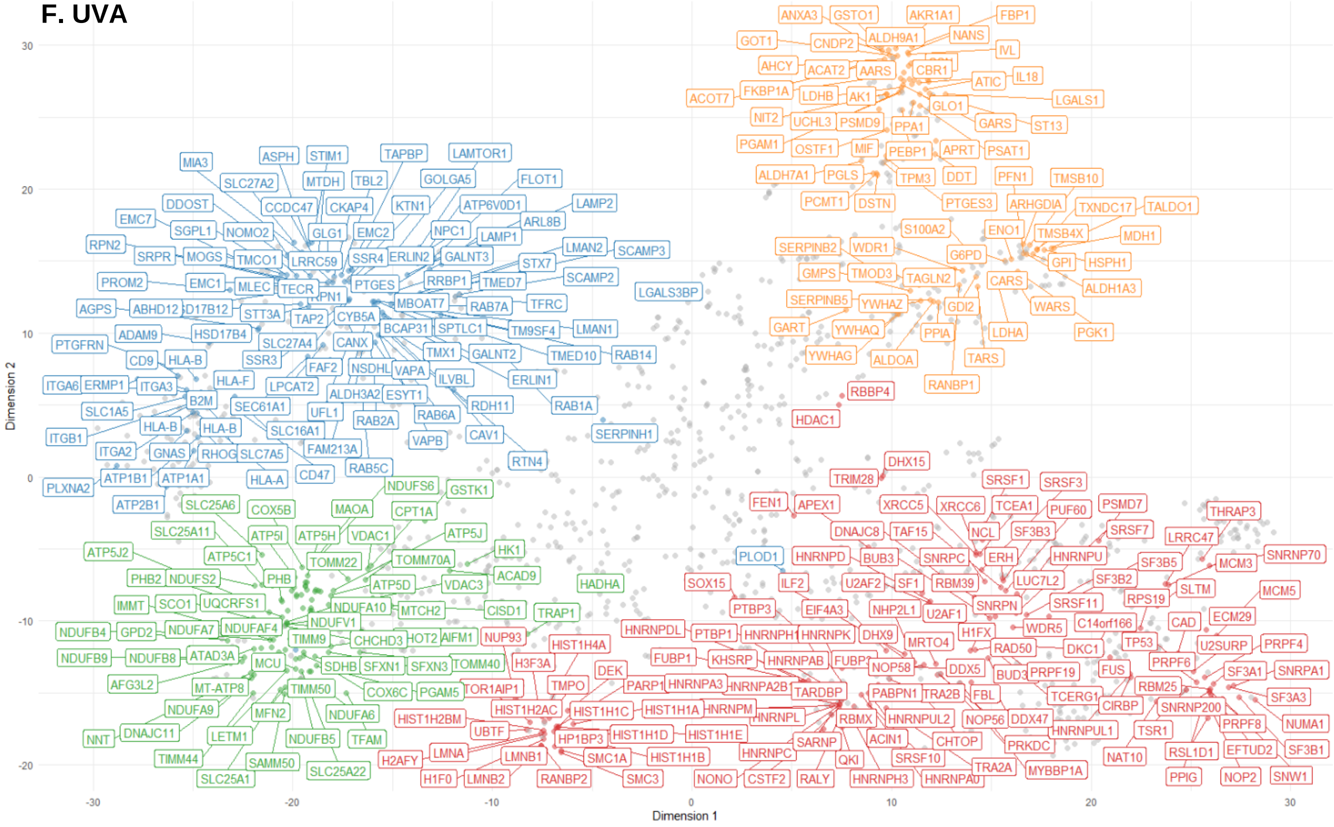


**Figure S1. Choice of imputation method based on t-SNE analysis. Related to Figures 1 and 2.**

**(A)** 2D t-SNE plot of all samples. The dataset was filtered for 50% of valid values in all the analysis performed in this paper. The remaining missing values were imputed as 0. The labels show the markers of subcellular localization used in this study. Red = nucleus, green = mitochondria, blue = secretory organelles and orange = cytosol. **(B)** 2D t-SNE plot of the whole dataset, considering imputation by Bayesian PCA Missing Value Estimation. **(C)** 2D t-SNE plot of the entire dataset, considering imputation by QRILC (Quantile Regression Imputation of Left-Censored data). **(D)** For proteins containing 4 valid values in a given fraction, the missing value was imputed as the average of the 4 valid values. The remaining missing values were imputed as the minimum value of the dataset. The figure shows the 2D t-SNE representation of this imputation method. **(E)** The subcellular map represents the 2D t-SNE plot of control samples under the imputation method of choice (average + minimum). **(F)** The subcellular map represents the 2D t-SNE plot of UVA samples under the imputation method of choice (average + minimum).

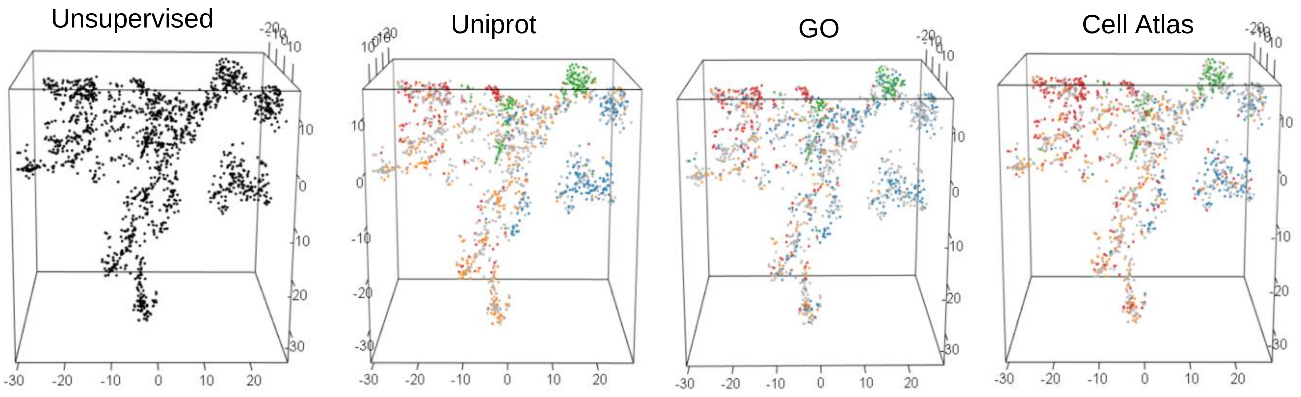


**Figure S2. Unsupervised and supervised 3D t-SNE analysis of the fractionation dataset. Related to Figure 2.** For supervised analysis, proteins were colored according to subcellular classifications from the Cell Atlas, Uniprot and Gene Ontology databases. All graphs were ploted with the same seed and same perplexity parameter of 30. Only proteins exclusively present in one of the 4 main compartments were colored in the analysis (Red = nucleus, orange = cytosol, blue = secretory organelles, green = mitochondria and gray = multilocalized proteins). As we were only inspecting data quality in this case, graphs were plotted for the entire dataset.

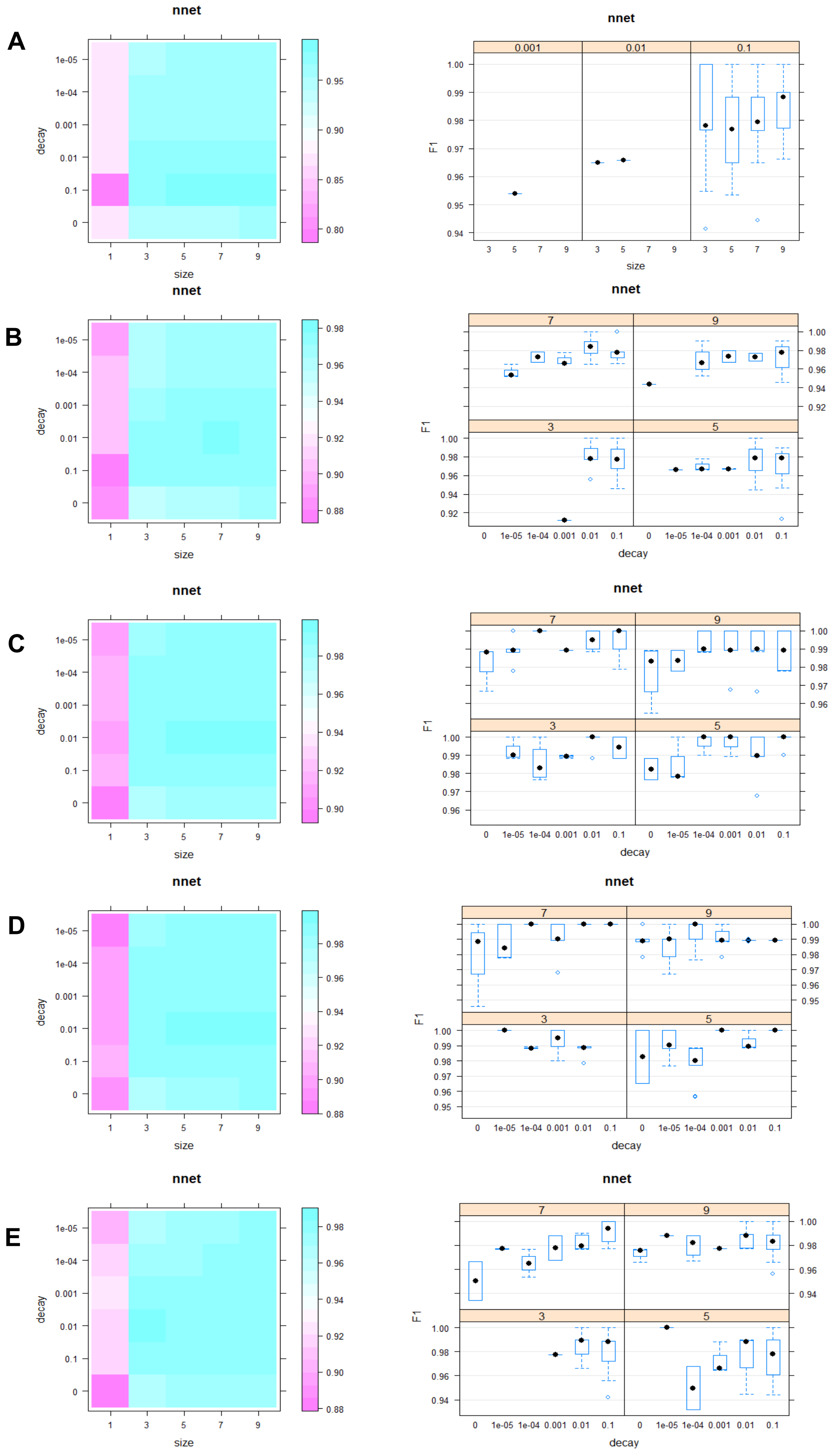

A
B
C
D
E
nnet
decay
size
F1
1e-05
1e-04
0.001
0.01
0.1
0
1
3
5
7
9

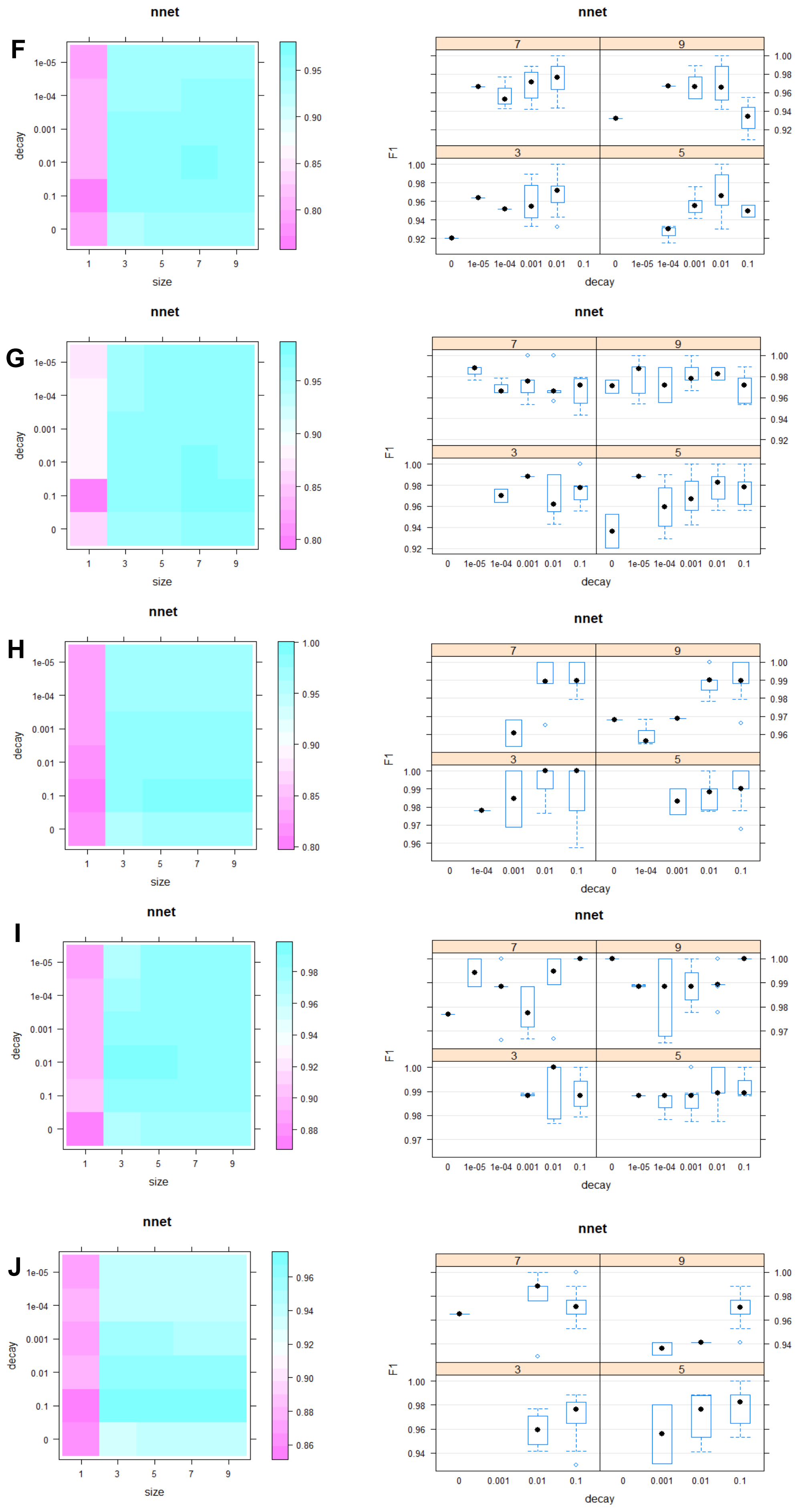

F
nnet
decay
size
nnet
F1
decay
G
nnet
decay
size
nnet
F1
decay
H
nnet
decay
size
nnet
F1
decay
I
nnet
decay
size
nnet
F1
decay
J
nnet
decay
size
nnet
F1
decay

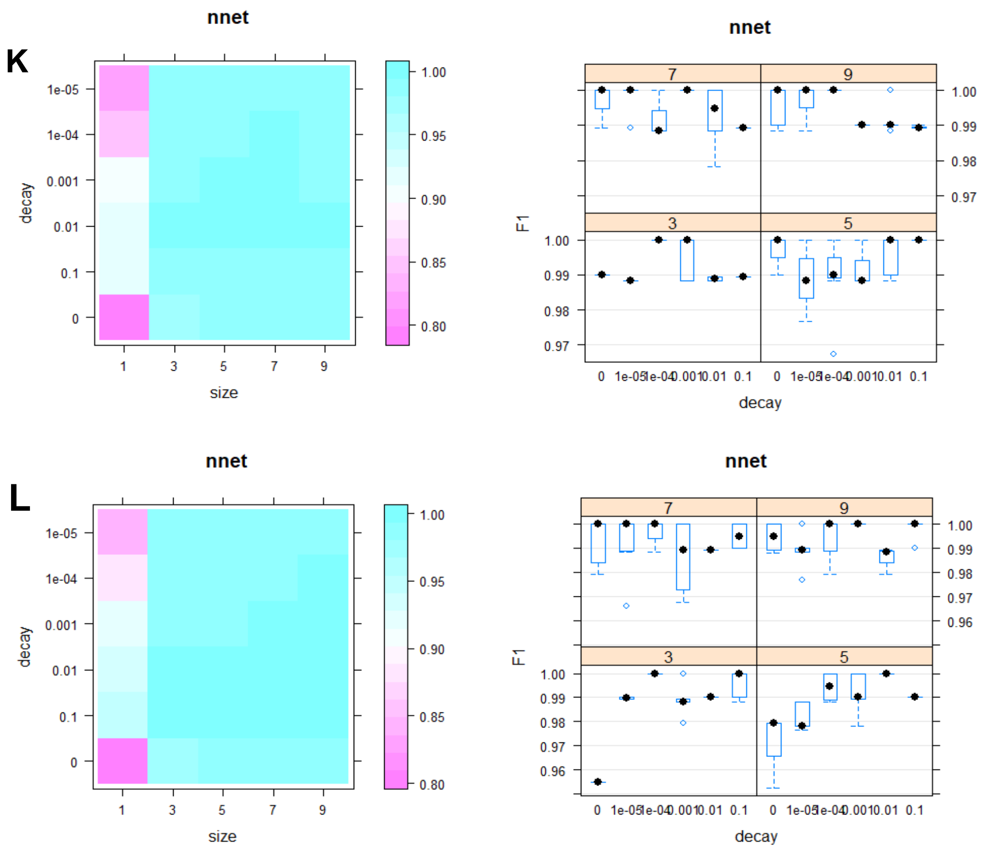


**M**

| Map | Decay | Size |
|---|---|---|
| Control: replicate 1 | 0.1 | 7 |
| Control: replicate 2 | 0.01 | 7 |
| Control: replicate 3 | 0.0001 | 3 |
| Control: replicate 4 | 0.0001 | 7 |
| Control: replicate 5 | 0.1 | 9 |
| UVA: replicate 1 | 0.01 | 9 |
| UVA: replicate 2 | 0.1 | 3 |
| UVA: replicate 3 | 0.1 | 3 |
| UVA: replicate 4 | 0.01 | 3 |
| UVA: replicate 5 | 0.1 | 7 |
| Consensus controls | 0.01 | 5 |
| Consensus UVA | 0.1 | 3 |

**Figure S3. Performance of the Averaged Neural Networks Classifier. Related to Figure 2.** Optimization of the hyperparameters of the classifier (size and decay). The F1 scorres were used as metrics to define the optimal hyperparameters. **(A - E)** Performance of the classifier across five folds of the training data, for each of the 5 replicates of control samples. **(F - J)** Performance of the classifier across five folds of the training data, for each of the 5 replicates of irradiated samples. **(K)** Performance of the classifier across five folds of the training data, for joint analysis of all controls. This analysis was performed to obtein a consensus classification for a given biological condition. **(L)** Performance of the classifier across five folds of the training data, for joint analysis of all irradiated samples. **(M)** Optimal hyperparameters chosen for the averaged neural networks analysis, considering both the joint analysis as well as the analysis of each replicate separately.

A

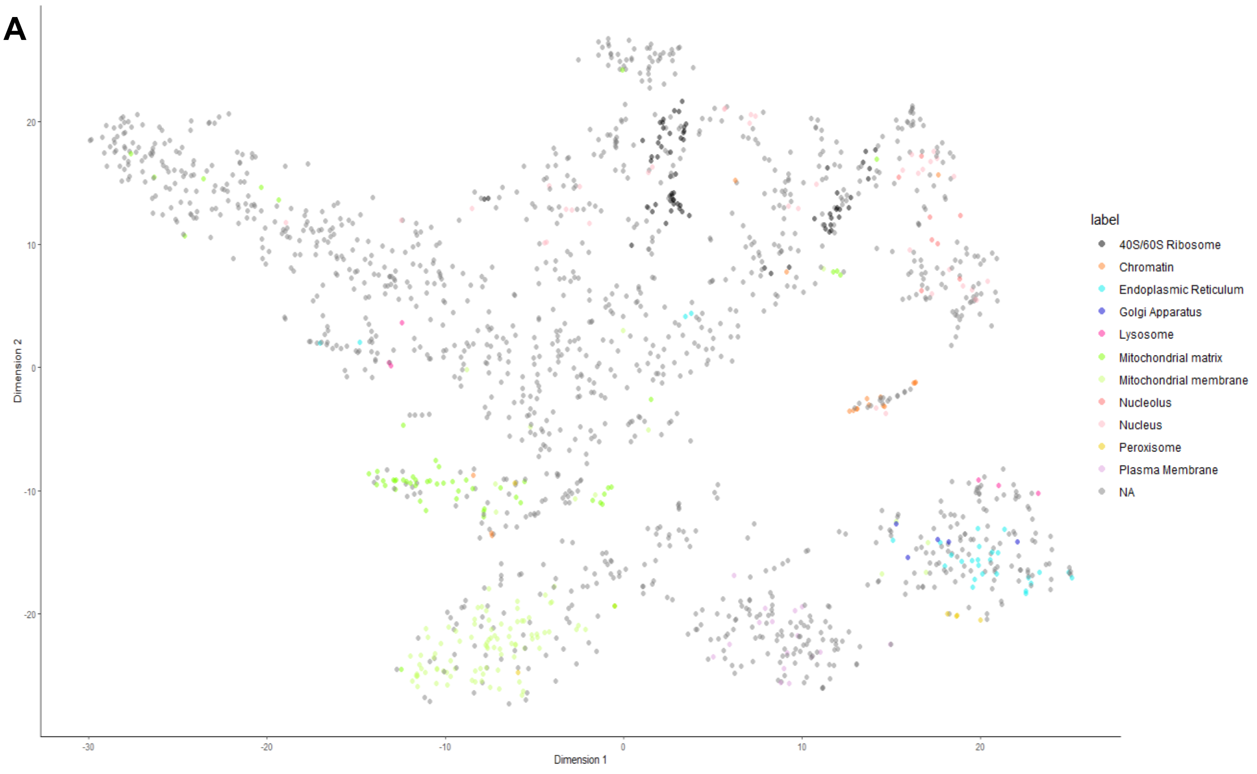

label
40S/60S Ribosome
Chromatin
Endoplasmic Reticulum
Golgi Apparatus
Lysosome
Mitochondrial matrix
Mitochondrial membrane
Nucleolus
Nucleus
Peroxisome
Plasma Membrane
NA
Dimension 1
Dimension 2


B

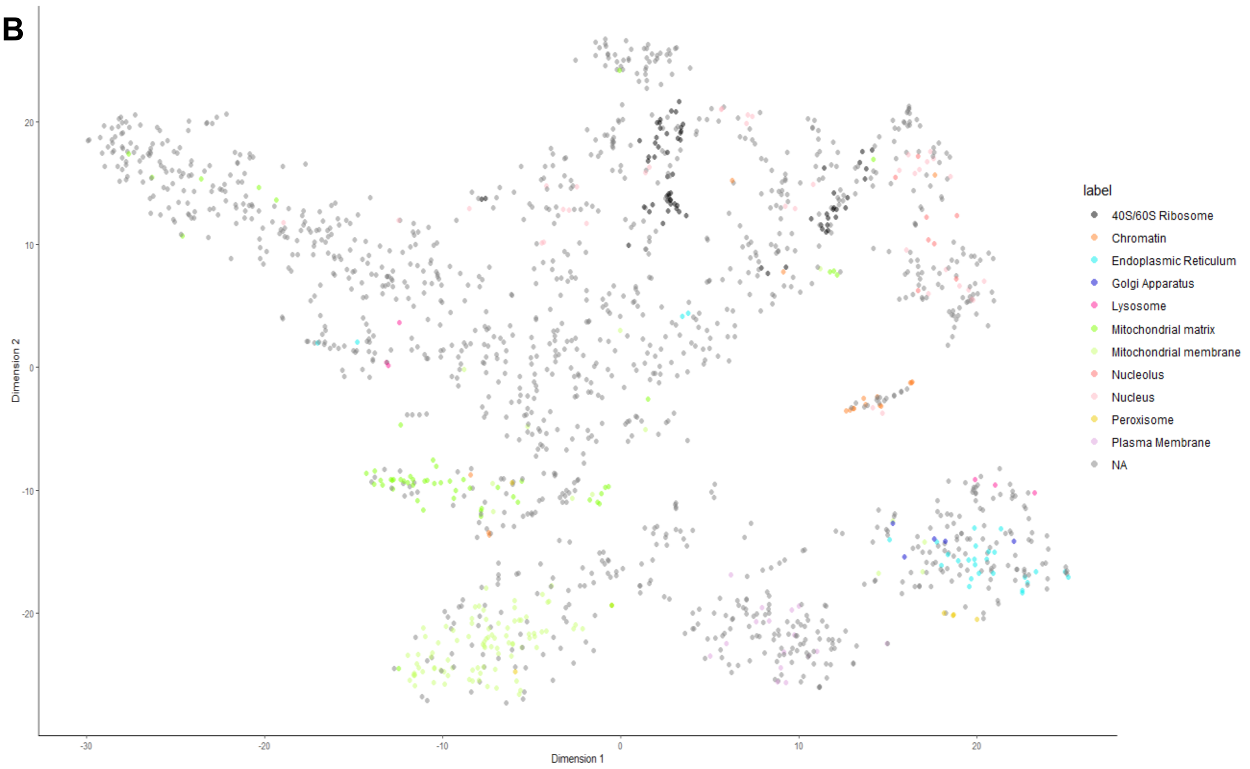

label
40S/60S Ribosome
Chromatin
Endoplasmic Reticulum
Golgi Apparatus
Lysosome
Mitochondrial matrix
Mitochondrial membrane
Nucleolus
Nucleus
Peroxisome
Plasma Membrane
NA
Dimension 1
Dimension 2

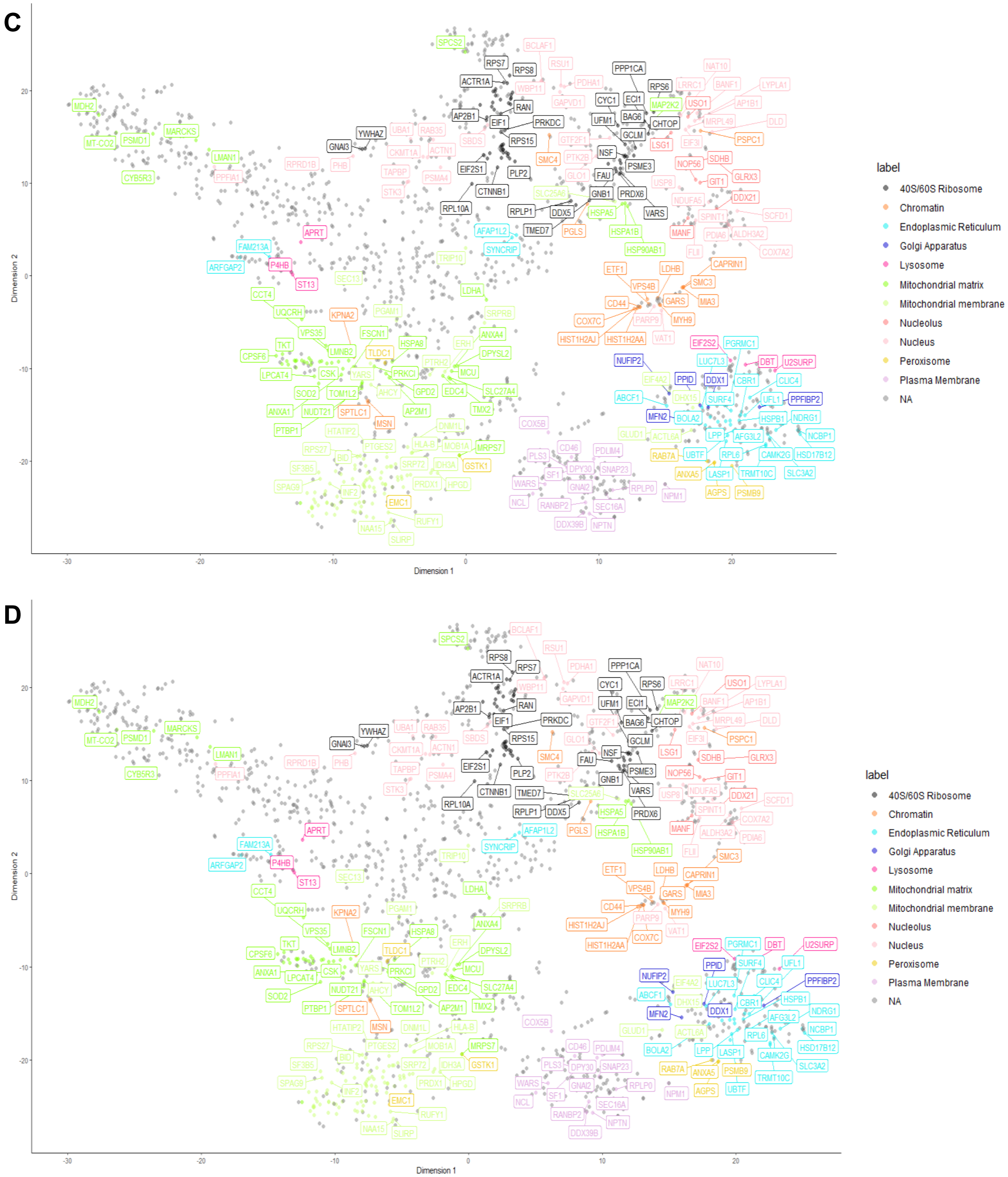


**Figure S4. 2D t-SNE plot showing through colors clusterization of sub-organellar niches, secretory organelles and protein complexes. Related to Figure 2. (A)** 2D t-SNE plot of control samples without label for gene names. **(B)** 2D t-SNE plot of irradiated samples without label for gene names. **(C)** 2D t-SNE plot of control samples with labels for gene names. **(D)** 2D t-SNE plot of irradiated samples with labels for gene names.

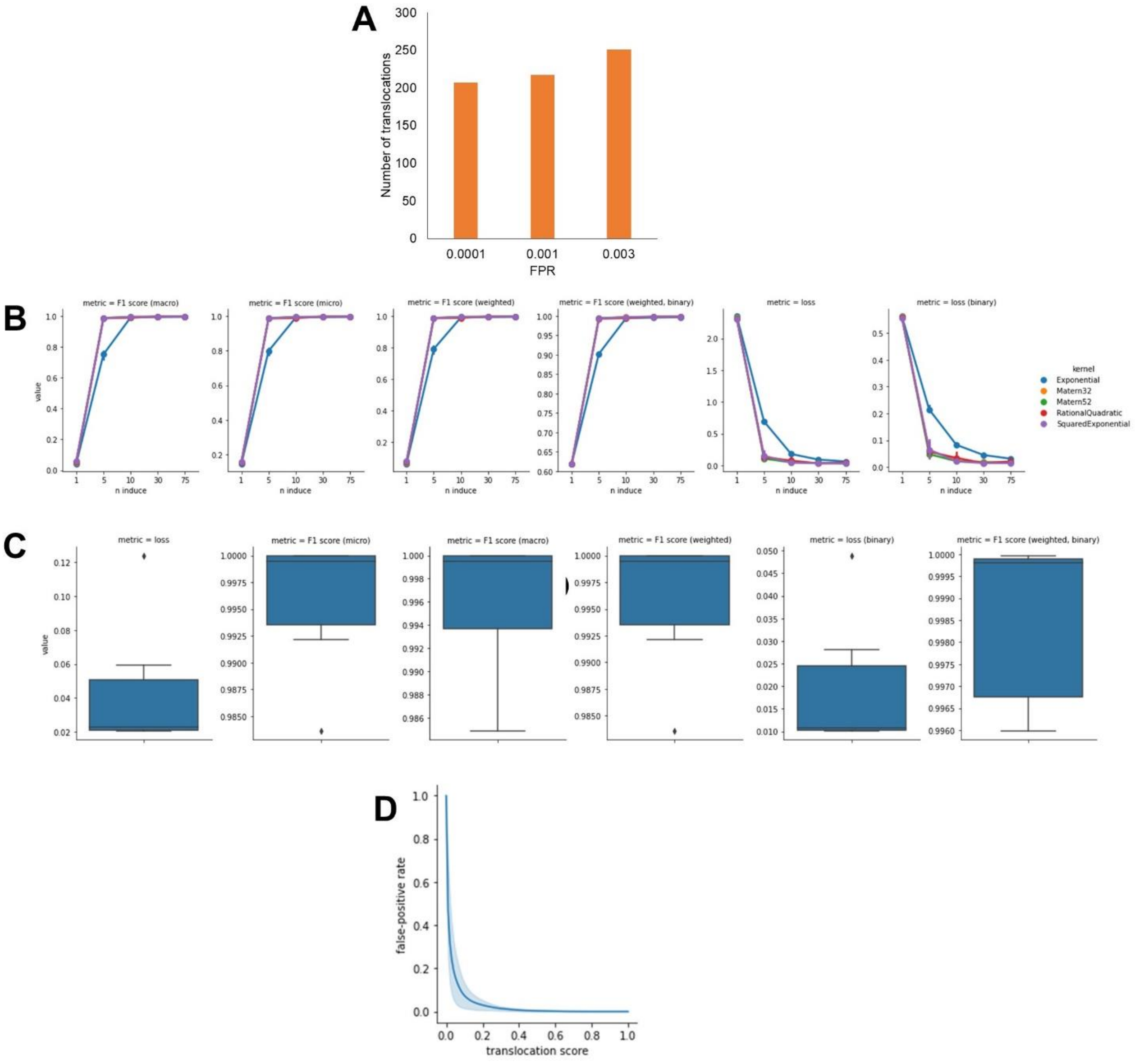


**Fig. S5. Performance of the Stochastic Variational Gaussian Process Classifier. Related to Figure 3.**

**(A)** Number of translocations identified with different FPR cutoff levels. **(B)** Optimization of the hyperparameters of the classifier (n induce and kernel type). The F1 scorres were used as metrics to define the optimal hyperparameters. The y axis represents the metrics as indicated in the titles of the plots. **(C)** Performance of the classifier across five folds of the training data. **(D)** False-positive rates for localization prediction based on the classifier performance of the test data.

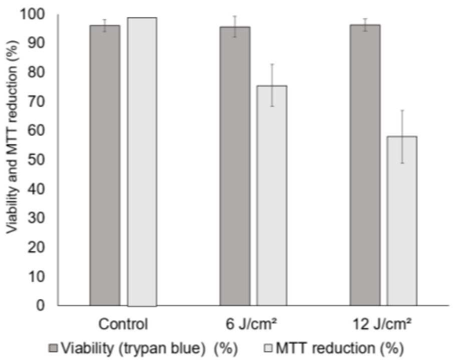


**Fig. S6. Cellular viability and reductive power upon UVA irradiation. Related to Figure 8.**
MTT and trypan blue assays were used to evaluate the cellular viability and reductive power of HaCaT cells under environmentally relevant UVA doses. Bars represent means and error bars represent standard deviation (n = 3). Percentage of MTT reduction is calculated as the ratio between the absorbance of formazan reduced by irradiated cells, and the absorbance of formazan reduced by control cells. Thus, the measurements of irradiated cells are relative to the measurements of controls, and per definition controls present 100% of MTT reduction capacity.

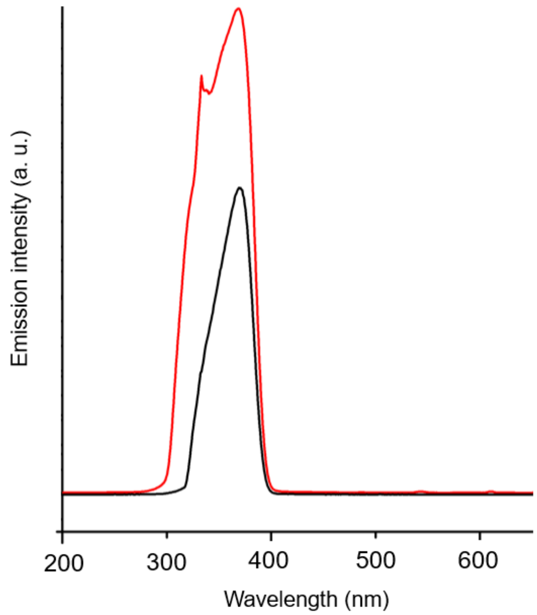


**Fig. S7. Emission spectra of the SOL-UV solar simulator equipped with a Xenon arc lamp and infrared bandpass blocking filter. Related to STAR Methods.** The spectra are shown with and without (in black and red respectively) the UVB-blocking filter.
